# Maximally predictive ensemble dynamics from data

**DOI:** 10.1101/2021.05.26.445816

**Authors:** Antonio C. Costa, Tosif Ahamed, David Jordan, Greg J. Stephens

## Abstract

We leverage the interplay between microscopic variability and macroscopic order to connect physical descriptions across scales directly from data, without underlying equations. We reconstruct a state space by concatenating measurements in time, building a maximum entropy partition of the resulting sequences, and choosing the sequence length to maximize predictive information. Trading non-linear trajectories for linear, ensemble evolution, we analyze reconstructed dynamics through transfer operators. The evolution is parameterized by a transition time *τ* : capturing the source entropy rate at small *τ* and revealing timescale separation with collective, coherent states through the operator spectrum at larger *τ*. Applicable to both deterministic and stochastic systems, we illustrate our approach through the Langevin dynamics of a particle in a double-well potential and the Lorenz system. Applied to the behavior of the nematode worm *C. elegans*, we derive a “run-and-pirouette” navigation strategy directly from posture dynamics. We demonstrate how sequences simulated from the ensemble evolution capture both fine scale posture dynamics and large scale effective diffusion in the worm’s centroid trajectories and introduce a top-down, operator-based clustering which reveals subtle subdivisions of the “run” behavior.

**POPULAR SUMMARY:** Complex structure is often composed from a limited set of relatively simple building blocks; such as novels from letters or proteins from amino acids. In musical composition, e.g., sounds and silences combine to form longer time scale structures; motifs form passages which in turn form movements. The challenge we address is how to identify collective variables which distinguish structures across such disparate time scales. We introduce a principled framework for learning effective descriptions directly from observations. Just as a musical piece transitions from one movement to the next, the collective dynamics we infer consists of transitions between macroscopic states, like jumps between metastable states in an effective potential landscape.

The statistics of these transitions are captured compactly by transfer operators. These operators play a central role, guiding the construction of maximally-predictive short-time states from incomplete measurements and identifying collective modes via eigenvalue decomposition. We demonstrate our analysis in both stochastic and deterministic systems, and with an application to the movement dynamics of an entire organism, unravelling new insight in long time scale behavioral states directly from measurements of posture dynamics. We can, in principle, also make connections to both longer or shorter timescales. Microscopically, postural dynamics result from the fine scale interactions of actin and myosin in the muscles, and from electrical impulses in the brain and nervous system. Macroscopically, behavioral dynamics may be extended to longer time scales, to moods or dispositions, including changes during aging, or over generations due to ecological or evolutionary adaptation. The generality of our approach provides opportunity for insights on long term dynamics within a wide variety of complex systems.

## INTRODUCTION

The constituents of the natural world combine to form a dizzying variety of emergent structures with effective laws across a wide range of scales. Sometimes these structures are obvious without referencing their underlying components; we needn’t, famously, link bulldozers to quarks [1]. But many are more ambiguous, especially in complex systems. How, for example, do we precisely identify behavioral states [2] or important patterns of collective neural activity [3]? When the scales are widely disparate, our best approach is often separate phenomenological modeling. In other cases however, a principled integration of fine-scale degrees of freedom can yield coarse-grained theories that successfully capture and predict large-scale structure. When we do make a connection, we are rewarded with greater insight than from either scale alone. We show here that it is possible to accomplish such coarse-graining directly from data.

Approaches from statistical physics, such as ensembles or kinetic theory (see e.g. [4]), provide important illustrations of successful cross-scale analysis. But powerful as they are, their implementation requires detailed knowledge of the underlying dynamics, symmetries and conservation laws, or an appropriate parameterization for formal methods like the renormalization group [5]. How can we emulate such success for systems for which we lack comparable understanding? One intriguing recent approach is to apply statistical inference to characterize how model selection, assessed through the Fisher information matrix, can systematically vary across scales [6, 7]. Another is to seek renormalization-type scaling [8, 9] from measurements.

Timescale separation offers a complimentary direction; coarse-grained degrees of freedom often vary slowly compared to their microscopic constituents. In the Langevin and Einstein approach to Brownian motion, for example, collisions experienced by a particle in a heat bath are treated as Gaussian white noise, allowing for the prediction of larger scale diffusive dynamics (see e.g. [10]). While timescale separations can be imposed, as in chemical kinetics when one assumes that some reactions are “fast” and essentially at equilibrium, we instead introduce an approach which allows such separations to be inferred, thus enabling the direct identification of slow dynamics. Slow degrees of freedom have been the focus of substantial previous work, from hydrodynamic Lyapunov modes in high dimensional dynamical systems [11] to neuroscience [12] and molecular dynamics [13].

We introduce a general framework for the principled extraction of coarse-grained, slow dynamics directly from time series data. Using known models, our first results illustrate this approach with the stochastic (finite-temperature) dynamics of a particle in a double-well potential and the deterministic chaos of the Lorenz system. To fully capture short-time dynamics we expand the dimensionality of the naive measurement space and partition the expanded space to maximize predictive information. We then coarse grain this maximally-predictive state space into metastable states in a way that preserves the Markovianity of the metastable dynamics. We thus simultaneously find an effective representation of both the states and of their dynamics. In doing so, we obtain a dynamics in which memory is subsumed into the representation, dramatically simplifying the resulting description. The ensemble dynamics are compactly represented as a transfer operator which encodes the time-evolution of state-space densities as a Markov transition matrix. These are generally known as Perron-Frobenius operators, of which a continuous analog is the Fokker-Planck equation [14].

For systems in which the fundamental equations are unknown, we focus on organism-scale behavior, where long timescales are of direct interest [15–18]. For behavior, navigation provides a relevant example; how do persistent states underlying search strategies emerge from much faster posture dynamics? An organism must successfully navigate its environment to improve its chances of feeding, mating, and survival, and substantial theoretical work has investigated the principles of environmental search, including chemotaxis, infotaxis and levy flights [19–21]. While environmental search is usually considered at the level of movement of the body centroid, differing modes of movement are generated by modulating neuromuscular control systems and their resulting postural dynamics. Human running and walking, for example, have discernibly different patterns of limb movement resulting in different locomotory speeds. From sequences of body posture it should be possible, in principle, to identify persistent states, as well as the behavioral strategies which modulate their frequency, thus linking distinct dynamics across scales.

Applied to the undulatory movement of the nematode *C. elegans* while foraging, we find that that “run and pirouette” kinetics naturally emerge from maximally-predictive ensemble dynamics of posture. We demonstrate that sequences simulated from the ensemble evolution capture both fine scale posture dynamics and large scale effective diffusion in the worm’s centroid trajectories. Finally, we introduce a top-down, operator-based clustering which reveals subtle subdivisions of the “run” behavior.

## THE TRANSFER OPERATOR PERSPECTIVE

We take a dynamical systems perspective but trade individual trajectories for ensembles, ultimately seeking long-lived dynamics in the patterns of stochastic state space transitions. The dynamics of ensembles is governed by operators which “transfer” the ensemble state from one time to another. An important advantage of this approach is that the dynamics are straightforwardly extracted as operator eigenvalues and eigenvectors.

We consider the general setting of the time evolution of a system with both deterministic and noisy influences through a set of differential equations for the evolution of the system’s state *x* ∈ℝ^*D*^. While such equations provide important information about the finescale properties of the dynamics, extracting longer time scale, coarse-grained properties from them is often challenging, indeed this is the primary challenge of kinetic theory (see e.g. [4]). Even when *D* is small, there are large scale dynamical patterns, such as unstable periodic orbits (UPOs) in chaotic dynamical systems [22], whose properties we cannot immediately infer from the knowledge of the equations of motion.

The motion of a particle trapped in a double-well potential at finite temperature provides an explanatory model dynamics, Fig. 1(A-left). If the barrier height is large compared to the temperature, the particle will hop between wells with almost vanishing probability, see e.g. [23]. Studying a single trajectory for only a short amount of time is misleading, as the particle mostly fluctuates around only one of the wells. If, however, we consider an *ensemble* of trajectories, or the time evolution of the probability densities, the two-well nature of the system becomes apparent, Fig. 1(A-middle). For both this example and Langevin dynamics in general, state space densities evolve according to a Fokker-Planck equation,

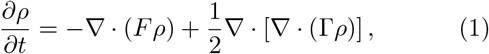

where *F* : ℝ^*D*^ →ℝ^*D*^ governs the deterministic dynamics, Γ = *γγ*^⊤^ and *γ* : ℝ^*D*^ →ℝ^*D*^ is the strength of the noise. By defining the linear operator ℒ such that 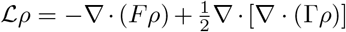, the ensemble evolution equation can be written as,

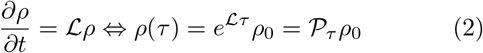

where *ρ*_0_ is an initial density, and 𝒫_*τ*_ is the Perron-Frobenius (PF) operator [24–27] that transfers the ensemble density from time *t* = 0 to a finite time *τ* in the future. We note that ℒ and therefore 𝒫_*τ*_ are not restricted to evolving dynamics of the Fokker-Planck class, extending equally to other stationary white noise processes and even deterministic dynamics [26–28], such as the Liouville operator of classical Hamiltonian dynamics. We leverage this universality to study stochastic and deterministic systems through a single framework.

**FIG. 1.**
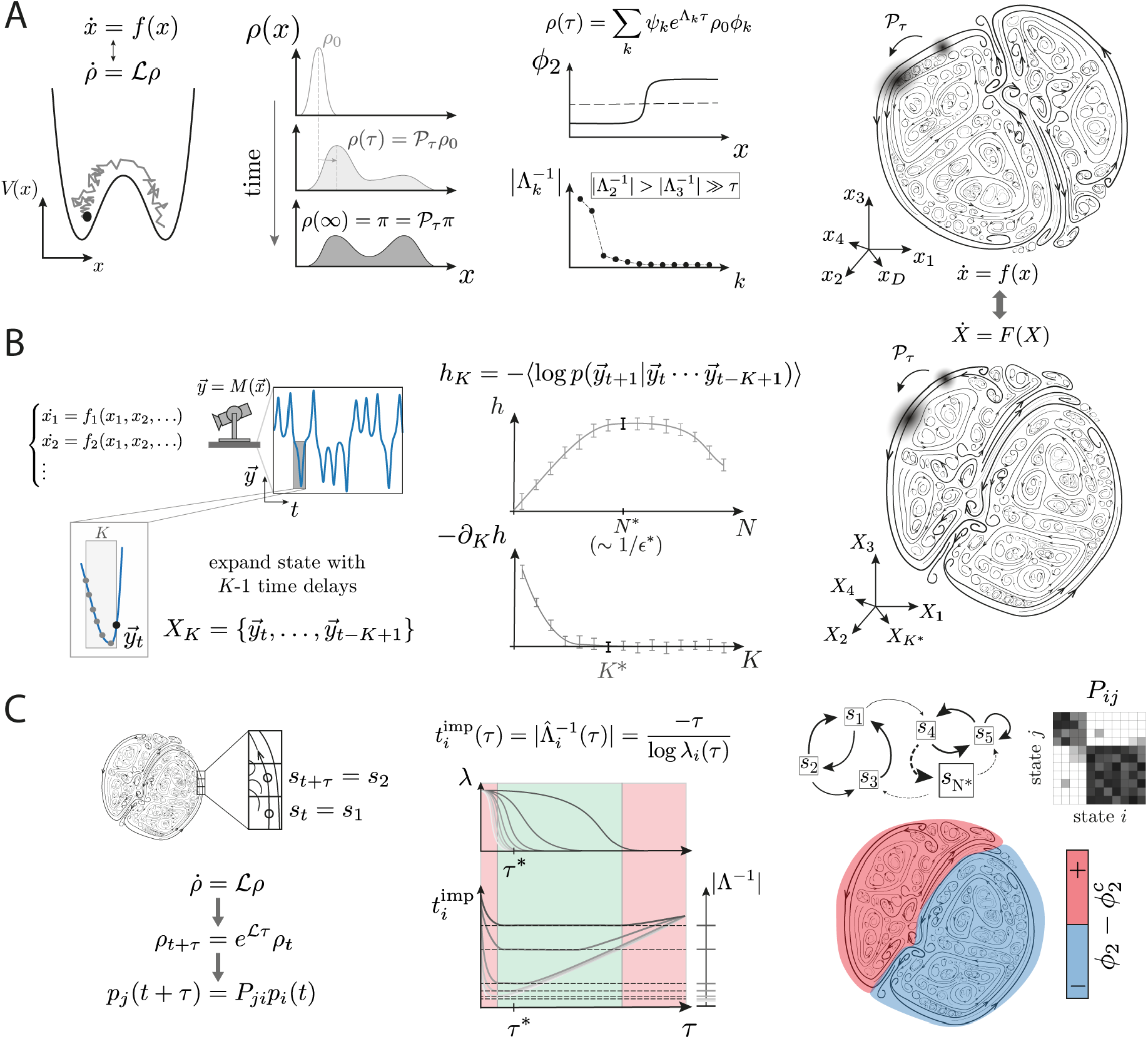
Maximizing predictive information for state space reconstruction and ensemble dynamics. (A) The ensemble approach naturally identifies large scale structures in systems with a large degree of unpredictability, for which individual short trajectories are less informative (left, particle in a double-well at finite-temperature). We trade non-linear dynamics in the state space 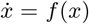 for equivalent linear dynamics in density space 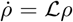. In discrete time, the ensemble evolution is dictated by the action of transfer operators 𝒫_*τ*_ = *e* ^ℒ *τ*^, evolving densities in time by a time scale *τ*. We achieve coarse-graining through the operator eigenfunctions and eigenvalues; the eigenfunction with Λ = 0 corresponds to the invariant distribution while the others decay on timescales 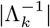. The eigenvalue gap is a principled guide to timescale separation: *ϕ*_2_ splits the invariant density into the two wells and 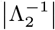 captures the hopping timescale. We leverage transfer operators to extract principled, coarse-grained descriptions of complex systems exhibiting intricate state-space structure (right). (B) We reconstruct the state space by concatenating measurements in time to maximize predictability. For each space of *K* delays we find the number of bins *N*\* that yields a maximum entropy partition and add delays until the entropy rate *h* is unchanging, *∂*_*K*_*h* (*K*\*) ≈ 0. Our reconstructed states *X* = {*y*_*t*_, …, *y*_*t*−*K*\*+1_} preserve the ergodic properties of the sampled system, including the density dynamics 𝒫_*τ*_. (C-left) We extract ensemble dynamics through the Perron-Frobenius operator 𝒫_*τ*_ approximated as the Markov transition matrix *P*_*ij*_ between state space partitions {*s*_*i*_} with transition time *τ*. *P*_*ij*_ provides a predictive model of ensemble dynamics applicable to both deterministic and stochastic systems. (C-middle) The inferred relaxation times of the underlying dynamics, *t*_imp_, implicitly depend on the choice of *τ*. For *τ* → 0, *t*_imp_ exhibit a short transient; *P*_*ij*_ is close to the identity matrix and the eigenvalues are nearly degenerate about 1 (left red shaded region). For *τ* ≫ *τ* * the eigenvalues are approximately constant as *P*_*ij*_ is determined primarily by fluctuations about the invariant distribution and relaxation times grow linearly (right red region). We choose *τ* * after the initial transient, when the regime in which the long-time dynamics are approximately Markovian and the spectral gap is the largest (green shaded region). (C-right) The eigenvectors *ϕ* of *P*_*ij*_ (*τ* *) naturally reveal coherent structures (red and blue), with the corresponding eigenvalues *λ* indicating the slow transitions between them.

Finding tractable analytic expressions of the PF operator is difficult in all but toy systems [23]. However, such operators can be effectively approximated by a Markov transition matrix between state space partitions [14, 27], and we exploit such an approximation to bridge the gap between fine scale and coarse-grained phenomenological descriptions of complex dynamical systems.

In particular, we focus on the operator eigenvalue spectrum, which, in the context of our Markov approximation, naturally orders the timescales of the system. In the two-well example, Fig. 1(A-middle), the equilibrium distribution *π* emerges as the eigenfunction with unit eigenvalue of 𝒫_*τ*_, 𝒫_*τ*_ *π* = *π*, while the exponentially-long timescale for hopping between wells is captured by the next-largest eigenvalue [23]. In general, the operator approach provides a hierarchy of timescales, which can naturally reveal a timescale separation and thus an effective phenomenological description of the system through the associated long-lived eigenfunctions. These eigenfunctions can capture important emergent coarsegrained properties of the dynamics such as transitions between multiple metastable states, Fig. 1(A-right). In the context of chemical kinetics, for example, such eigen-fuctions provide ideal reaction coordinates or order parameters [13, 29, 30], which are analogous to the committor function of transition path theory [30–32].

## PREDICTIVE INFORMATION FOR STATE SPACE RECONSTRUCTION

In real-world complex systems, individual measurements, even if high-dimensional, rarely capture the full set of variables that constitute the state space. Analyses based solely on such measurements thus ignore important fine-scale information. For example, the state space for a simple pendulum is two dimensional, consisting of the angular position and angular velocity. If we attempt to reconstruct the dynamics from the position alone, we lack important information about whether the pendulum is on an upswing or a downswing, and this will manifest as predictive information we have missed. We can recover this information using a time delay embedding [33–37], as we demonstrate in our model system examples. We show here, in general, how to construct a state space that contains maximal predictive information.

Consider a set of incomplete measurements of an unknown dynamical system, 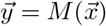, where *M* is a measurement function mapping the underlying state space 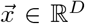 into our measurements 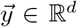, for which typically *d* < *D*, Fig. 1(B-left). We expand the putative state by adding *K* − 1 time delays to the measurement time series, sampled on a time scale *δt*, yielding a candidate state space *X*_*K*_ ∈ ℝ^*d*×*K*^. We quantitatively characterize the unpredictability of the candidate state space through the entropy rate of the symbolic dynamics resulting from partitioning each *K × d*-dimensional putative state space into *N* Voronoi cells through clustering (Methods). With a partitioning, the reconstructed dynamics are encoded as a row-stochastic transition probability matrix *P* which evolves a state-space density *p* by a time *δt*,

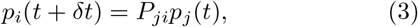

where we sum over repeated indices (Einstein’s summation convention). The entropy rate of the source is approached by estimating the entropy rate of the associated Markov chain for increasing values of *K*,

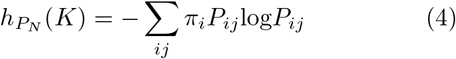

where *π* is the estimated stationary distribution of the Markov chain *P* ^1^. The Markov approximation of the entropies provides an estimate of the conditional entropies between discrete states *s*, ⟨−log [*p*(*s*_*j*_|*s*_*i*_)])⟩, where *i, j* ∈ {1, …, *N*}. Each discrete state contains a population of delay vectors 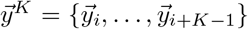, and therefore the entropy of the Markov chain provides an estimate of the sequence entropy of the time series,

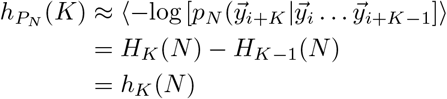

where *H*_*K*_(*N*) is the entropy of the *K*-gram symbolic sequence built by discretizing the 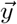 space into *N* partitions. Note that the entropy rate is a non-decreasing function on the number of partitions, N, which is equivalent to a typical state-space scale *ϵ*∼ 1*/N*. Importantly, we seek to preserve as much information as possible in the discretized state space, setting the number of partitions as the largest *N* after which the entropy rate stops increasing due to finite-size effects Fig. 1(B-middle). This yields the maximum entropy rate with respect to the number of partitions and thus approximates the Kolmogorov-Sinai (KS) limit of the entropy rate [38, 39]. Besides the KS entropy, other quantities such as information or correlation dimensions can in principle be obtained by studying the scaling of the measure with the partition size [38, 40, 41].

With increasing K, we approach the partition entropy rate of the source from above so that the difference,

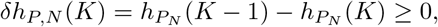

is a non-increasing function of *K. δh*_*P,N*_ (*K*) has been used to define measures of forecasting complexity in dynamical systems [42] and is the amount of information that has to be kept in the *K* − 1 time delays for an accurate forecast of the next time step. When *K* is too short, 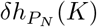 will be large, meaning that a large amount of information is required in order to make an accurate prediction. In systems with finite range correlations, there is a *K*\* for which *δh*_*P,N*_ (*K*\*) = 0, in which *K*\* corresponds to the amount of memory sitting in the measurement time series. Our state space reconstruction seeks *K*\* such that 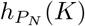 is minimized, i.e.,

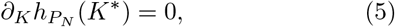

which corresponds to maximizing the predictive information. We recall the general definition of predictive information [43], 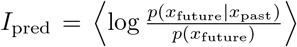. Defining *x*_past_ as the first *K* − 1 time steps in the time series and *x*_future_ as the *K*th time step,

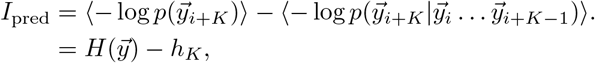

and thus with respect to a partition into *N* states, we have

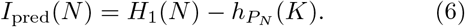

*I*_pred_(*N*) thus quantifies the reduction in the topological entropy of the measurement time series that results from the knowledge of the transition probabilities. The predictive information is maximized when 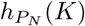 is minimized, which for a system with finite correlations is attained when 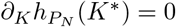.

Given a reconstructed state *X*_*K*\*_, the transition matrix *P*_*ij*_ (defined with timestep *τ* = *δt*), is an example of an approximated *transfer operator* 𝒫_*τ*_, Eq. (2) [14], Fig. 1(B-right). The transfer operator dynamics solely depends on the topology of the state space trajectories, which are guaranteed to be preserved by a state space embedding [34, 37]. In that sense, 𝒫_*τ*_ is in principle exactly preserved, and maintains all the properties of the underlying dynamics. In contrast, the set of non-linear equations of motion driving the dynamics of the reconstructed state 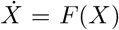, which are for instance required to obtain estimates of the local Jacobian, strongly depend on the geometric properties of the space (such as dimensions, metric, etc.), making it non-trivial to accurately approximate the underlying dynamics 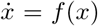, Fig. 1(A,B-right). The transfer operator formalism is therefore complementary to trajectory based approaches, providing a means to study large scale properties of the dynamics while being robust to the precise geometric properties of the reconstructed state.

## EXTRACTING COARSE-GRAINED DYNAMICS

In the state space reconstruction we encode ensemble dynamics in a matrix *P*_*ij*_ constructed by counting transitions between cells *A*_*i*_ and *A*_*j*_ in a transition time *τ* : formally this is a discrete, finite-time approximation to the transfer operator of Eq. (2). The eigenvalues of *P*_*ij*_ can directly reveal time-scale separation and the corresponding eigenvectors can be used to identify regions of state space where the system lingers, these are “macroscopic” metastable states. The structure of these regions and the kinetics between them offer a principled coarse-graining of the original dynamics. This coarse-graining is not imposed but rather follows directly from the ensemble dynamics, an important asset when analyzing complex systems.

### Choosing a transition time

Recall that the instantaneous ensemble dynamics is characterized by the generator ℒ of Eq. (2), Fig. 1(C-left). However, when working directly from measurements sampled at discrete times, the finite-time transfer operator 𝒫_*τ*_ is more immediately available. The operators ℒ and 𝒫_*τ*_ share the same set of eigenfunctions, while the eigenvalues 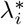 of 𝒫_*τ*_ are exponential functions of the generator eigenvalues 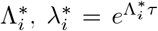. When the estimated eigenvalues *λ*_*i*_ are real, the relaxation of the inferred density dynamics is characterized by the implied relaxation times corresponding to each eigenvalue,

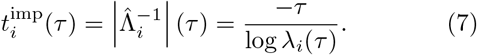

𝒫_*τ*_ lives in an infinite dimensional functional space, which we discretize using *N* basis functions: in our partitioned space, the basis functions are characteristic functions and the measure is piecewise constant (Methods). The truncation at finite *N* erases fine-scale information within the partition, and so the ability of the transfer operator approximation *P*_*ij*_ to capture long-time dynamics depends on the transition time *τ*, a parameter that we vary. For the state space reconstruction described in the previous section, we chose *τ* as the sampling time *τ* = *δt* in order to maximize short-time predictability and to capture the Kolmogorov-Sinai limit of the entropy rate of the dynamics, *ϵ* ∼ 1*/N* → 0, *δt* → 0 [39]. However, to accurately capture longer-time dynamics and metastable states, we vary *τ* and study changes in the inferred spectrum, Fig. 1(C-middle).

For *τ* →0, *P*_*ij*_ is close to an identity matrix (the system has little time to leave the current partition) and all eigenvalues are nearly degenerate and close to unity, Fig. 1(C-middle). For *τ* much longer than the mixing time of the system, the eigenvalues start collapsing and in the limit of *τ* → ∞ ⇒ *P*_*ij*_(*τ*) ∼ *π*_*j*_: the probability of two subsequent states separated by such *τ* becomes independent, the eigenvalues of *P*_*ij*_(*τ*) stop changing and *t*_imp_ grows linearly with *τ* for all eigenfunctions. In this regime, the transition probability matrix contains noisy copies of the invariant density akin to a shuffle of the symbolic sequence. This yields an effective noise floor, which is observed earlier (shorter *τ*) for faster decaying eigenfunctions. For intermediate *τ* there is a spectral gap, indicating that the fast dynamics have relaxed and we can isolate the slow dynamics. For such *τ*, the inferred relaxation times reach a constant value that matches the underlying long time scale 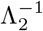, indicating that such dynamics are Markovian: the Chapman-Kolmogorov equation (see e.g. [44]) is verified 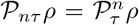, and thus 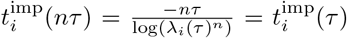^2^. Notably however, the dynamics on the discretized state space is not generally Markovian on all time scales since the discretization erases fine scale information. Nonetheless, a Markov model can approximate the slow dynamics arbitrarily well, provided that the discretization can be made arbitrarily fine and the transition time scale *τ* sufficiently long [45, 46]. We choose *τ* * after the initial transient, which corresponds to the regime in which the spectral gap is the largest and timescale separation is evident (green shaded region, Fig. 1(C-middle)).

### Identifying metastable states

Within our reconstructed ensemble evolution, coarse-grained dynamics can be identified between state space regions where the system becomes temporarily trapped, these are metastable states or almost invariant sets [47, 48]. We search for collections of states that move coherently with the flow: their relative future states belong to the same macroscopic set. Since the slowest decay to the stationary distribution is captured by the first non-trivial eigenfunction of the transfer operator, we search for the state space subdivision along this eigenfunction that maximizes the coherence of the resulting macroscopic sets, a previously introduced heuristic [47].

A set *S* is coherent when a state is more likely to remain within the set that it is to leave it within a time *τ*. We quantify this intuition through 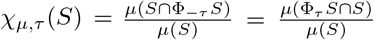, where *μ* is the invariant measure preserved by the invertible flow Φ_*τ*_. Given an inferred transfer operator *P*_*ij*_(*τ*) and its associated stationary eigenvector *π*, we can immediately compute *χ* (Methods),

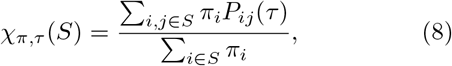

and we identify a coherent set when *χ*_*μ,τ*_ (*S*) is large. The approximation of transfer operators for the extraction of long-lived states has been successfully applied in a number of systems: from identifying metastable protein conformations in molecular dynamics simulations [13], to determining coherent structures in oceanic flows [28, 49, 50] (for reviews, see [14, 32, 51–54]).

To identify optimally-coherent metastable states through spectral analysis, we define a time-symmetric (reversibilized) transfer operator,

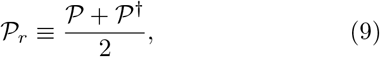

where 𝒫^†^ is the dual operator to 𝒫, pulling the dynamics backward in time; see Methods for the discrete numerical approximation, *P*_*r*_. While transfer operators describing ensemble dynamics are not generally symmetric [47, 55], the definition of coherence is invariant under time reversal: since the measure is time invariant, it does not matter in which direction we look for mass loss from a set [32, 56]. Besides the invariance property, an important benefit of working with *P*_*r*_ is that its second eigenvector *ϕ*_2_ provides an *optimal* subdivision of the state space into almost invariant sets, as shown in [56]. Given *τ* * and the corresponding *P*_*r*_(*τ* *), we identify metastable sets by choosing a subdivision along *ϕ*_2_ that maximizes an overall measure of coherence,

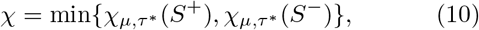

where {*S*^+^, *S*^−^} result from a partition at 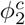: the optimal almost invariant sets are identified with respect to the sign of 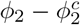, Fig. 1(C-right).

## RESULTS

### Brownian particle in a double well potential

We consider the Langevin dynamics of a particle in a double well potential *V* (*x*) = (*x*^2^ − 1)^2^ in thermal equilibrium, Fig. 2(A-left). The equation of motion for the position *x* of the particle is

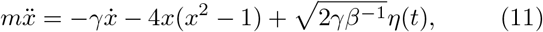

**FIG. 2.**
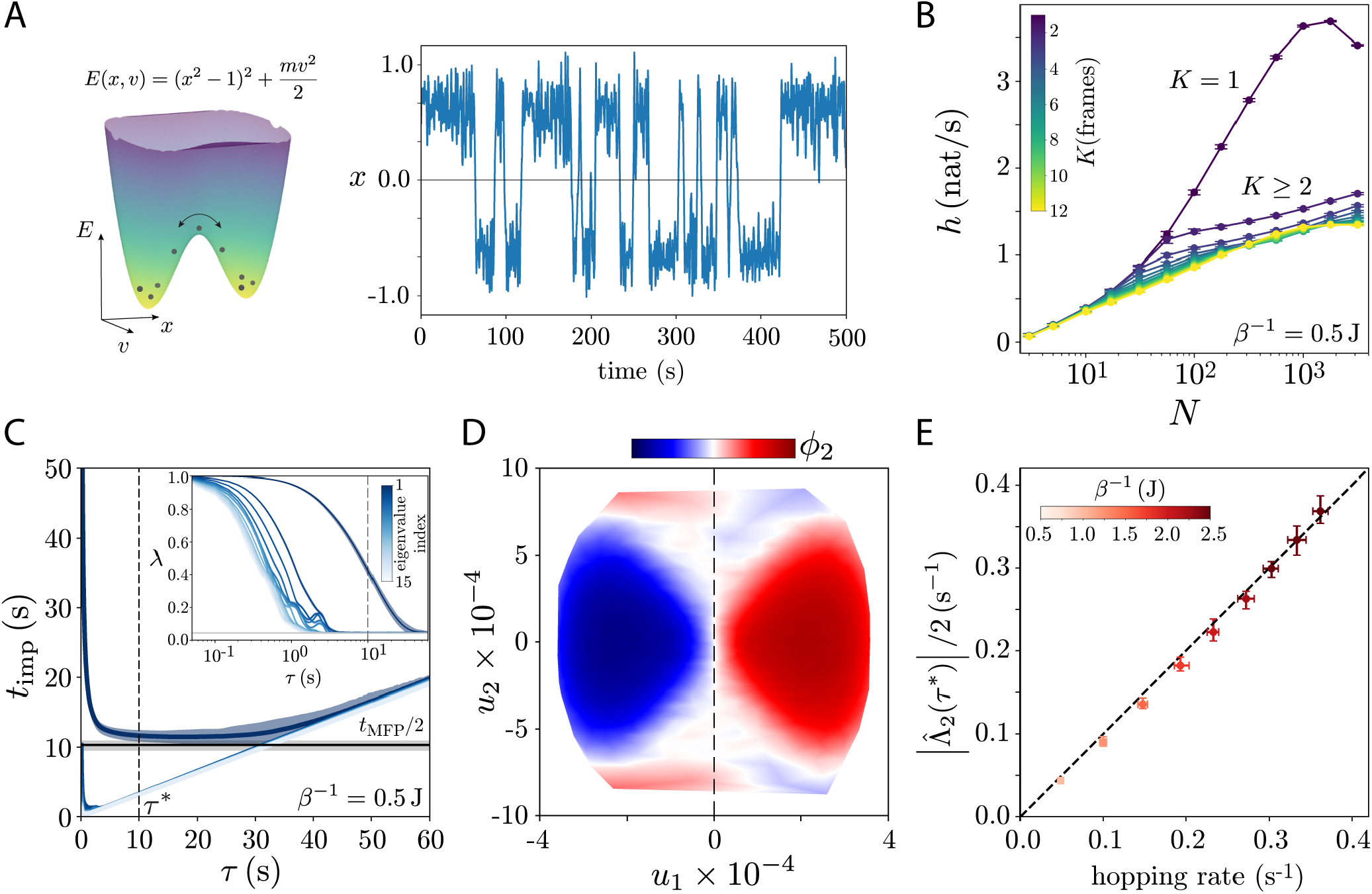
Maximally predictive ensemble dynamics of a Brownian particle in a double-well potential with viscous damping. (A) We use Langevin dynamics Eq. (12) to simulate a long (10^6^ s) trajectory of an underdamped particle in a double-well potential (left). To emulate the challenge of unknown systems, we measure only the position *x*(*t*) and show a sample of the trajectory with effective temperature *β*^−1^ = 0.5 J (right). (B) We concatenate the measurement with *K* delays, partition the resulting state with *N* partitions and show the entropy rate as a function of *K* and *N*. The behavior of *h*(*K, N*) reveals important information. The entropy rate changes dramatically as missing (and predictive) velocity information is incorporated into the state (*K >* 2 frames). Notably, the reconstructed state is not fully Markovian at *K* = 2 frames, a memory effect induced by the Euler-scheme used for simulation [57, 59, 60]. As expected for this continuous stochastic process *h*(*K, N*) → ∞ with increasing *N*, except when finite size effects result in the underestimation of the entropy rate. Furthermore, once the additional velocity information is incorporated, we see the approximate logarithmic divergence with *N* as is expected for a Markov-chain. We choose *K*\* = 7 frames and *N* = 1000 partitions to construct the finite-time ensemble evolution operator *P* (*τ*) and its reversibilized counterpart *P*_*r*_ (*τ*). (C) We plot the eigenvalues *λ*(*τ*) of *P*_*r*_ (*τ*) (inset) and corresponding implied timescale *t*_imp_(*τ*) for *β*^−1^ = *k*_*B*_*T* = 0.5 J. We choose a transition timescale *τ* * such that there is a clear separation; Markov dynamics in the first non-trivial eigenfunction *ϕ*_2_ are evident when *t*_imp_ is constant (dark blue). The short-*τ* transient results from a nearly diagonal transition matrix reflected in the accumulation of eigenvalues near 1. For large *τ* the eigenvalues stop changing as the sequence probabilities reach the noise floor (gray horizontal line) and become independent: in this regime *t*_imp_ grows linearly. The longest relaxation time is approximately constant after *τ* ∼ 5 s, converging to half the mean first passage time *t*_MFP_*/*2 (black horizontal line) as expected from theory. We choose *τ*\* = 10 s for subsequent analysis. (D) Contour plot of *ϕ*_2_ projected onto the (*u*_1_, *u*_2_) space for *β*^−1^ = 0.5 J. The sign of *ϕ*_2_ effectively splits the reconstructed state space into the two wells: *ϕ*_2_ *<* 0 corresponds to the right well, while *ϕ*_2_ *>* 0 corresponds to the left well. Note also that the high velocity regions | *u*_2_ | correspond to transition regions. (E) 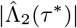 predicts the hopping rate across a range of temperatures. We choose *τ* * separately for each temperature to to reflect the different dynamics. The behavior of the longest implied time scale is qualitatively similar across temperatures, reaching a minimum at the plateau region, Fig. S2(B). We thus choose *τ* * as the time scale that minimizes the longest *t*_imp_. Error bars represents 95% confidence intervals over 200 non-overlapping trajectory segments of length 50000 s.

where *β*^−1^ = *k*_*B*_*T, m, γ, k*_*B*_ and *T* are the mass, damping coefficient, Boltzmann constant and temperature, respectively, and *η*(*t*) is a Gaussian white random variable capturing the random collisions with the heat bath. The dynamics is second order in the position *x*, and so we introduce the velocity 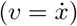 degree of freedom to rewrite Eq. (11) as a system of first order Ito stochastic differential equations evolving the state (*x, v*) in time (we use *m* = *γ* = 1 throughout),

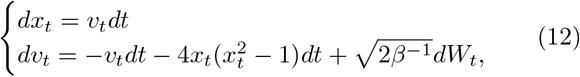

where *dW*_*t*_ is a Wiener process. The dynamics of (*x, v*) is first order and therefore fundamentally Markovian and memoryless. To emulate a real data example for which only a subset of the degrees of freedom is available, we measure only the position *x* of the particle, and seek to reconstruct the state space by adding time delays.

We generate long simulations of 10^6^ s sampled at *δt* = 0.05 s, with temperatures ranging between *β*^−1^ = 0.5 J and *β*^−1^ = 2.5 J (Methods): an example trajectory for *β*^−1^ = 0.5 J is shown in Fig. 2(A-right). We concatenate the *x* measurement with *K* delays, partition the resulting state with *N* partitions and show the entropy rate as a function of increasing *K* and *N*, Fig. 2(B). Notably, the behavior of *h*(*K, N*) reveals fundamental information about the properties of the measurement time series [39]. Since the generating equations are a continuous-time stochastic process, lim_*N*→∞_ *h*(*K, N*) = ∞. In addition, the rate of approach to infinity depends on the complexity of the random process [39]: for a Markov-chain approximation of the Langevin dynamics we expect *h ∝* log(*N*), as observed for *K* ≳ 2 frames, a result that we derive from the Cohen-Procaccia *h*(*ϵ, τ*) estimate of the entropy translated to our partition-based approach assuming *ϵ* ∼ 1*/N* (Eq. 3.71 in [39] for fixed *τ*). For small *N*, there are not enough partitions to see the difference between candidate state spaces. For large *N*, finite-sampling effects produce an underestimation of the entropy [38, 39], a drop we see most visibly for *K* = 1 frame. At intermediate *N*, where estimation effects are negligible, there is an abrupt change in the entropy rate from *K* = 1 to *K* > 2 frames = 0.1 s, indicative of the finite memory sitting in the measurement time series from the missing degree of freedom. Notably, the dynamics is not fully memoryless with *K* = 2 delays, as one would naively expect, a result due to the Euler-scheme update which induces memory effects on the infinitesimal propagator *P* (*x*_*k*+1_ *x*_*k*_, *x*_*k*−1_) [57–60]. The behavior of the entropy with *K* and *N* is qualitatively conserved across the range of temperatures studied here, Figs. S1(A),S5(A), and we choose *K*\* = 7 frames = 0.35 s for subsequent analysis. Our reconstructed state recovers the equipartition theorem, showing that momentum information is accurately captured by the reconstructed state, Fig. S1(B).

Using the reconstructed state space with *K*\* = 7 frames, we approximate the transfer operator by making a Voronoi tesselation through k-means clustering (Methods), and counting the transition probabilities between discrete states as a function of transition time scale *τ*. We construct the reversibilized transition matrix *P*_*r*_ and show the 15 slowest inferred relaxation times for *N* = 1000 partitions and transition times *τ*, Fig. 2(C). The longest relaxation time decays as the system thermalizes, and reaches its asymptotic limit after *τ* ∼ 5 s. As we increase *τ* to a few multiples of the hopping time scale, the eigenvalues stop changing and *t*_imp_ simply grows linearly with *τ* ; faster decaying eigenfunctions exhibit this behavior earlier.

To gain further intuition into the meaning of the longest implied timescale, we simulate the ensemble dynamics using the inferred transfer operator *P*_*ij*_ for *β*^−1^ = 0.5 J. In Fig. S2(A) we show the decay of second eigenfunction as a function of time and the coordinates (*u*_1_, *u*_2_), which are projections along the first two singular vectors of the state space embedding that match the position and velocity degrees of freedom, Fig. S1(C). We initiate the system with the density *ρ* concentrated in one of the *N* = 1000 partitions and use *τ* * = 10 s as the transition time. After *t* = 10 s the ensemble has distributed itself in one of the wells and at *t* = 40 s, it has almost completely relaxed to the invariant density, which here corresponds to the Boltzmann distribution.

The state space structure of the second eigenvector of the reversibilized transition matrix *ϕ*_2_ reveals a collective coordinate capturing transitions between the wells. In Fig. 2(D) we show a contour plot of *ϕ*_2_ in the (*u*_1_, *u*_2_) space for *β*^−1^ = 0.5 J where it is apparent that the sign of the eigenvector effectively splits the reconstructed state space into the two wells of the system. In fact, the maximum of *χ*, Eq. (10), is obtained at 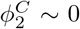 across temperatures, Fig. S2(C). Notably, due to the underdamped nature of the dynamics the separation between potential wells includes the high velocity transition regions |*u*_2_|, as expected from theory [61]. Indeed, 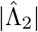 provides an excellent fit to the hopping rate which we estimate empirically by fitting an exponential function to the dwell time distributions in both wells, Fig. 2(E). We note that in order to examine different temperatures the choice of *τ* * should change accordingly to reflect the different resulting dynamics. Increasing the temperature reduces the hopping rates and therefore the amount of time it takes before the system relaxes to the stationary distribution. Nonetheless, the qualitative behavior of the longest inferred implied relaxation times is conserved across temperatures, Fig. S2(B): a short *τ* transient quickly converges to the time scale of hopping between wells, before the system completely mixes and the implied time scales grow linearly. To choose *τ* * consistently across temperatures we find the transition time that minimizes the longest reversibilized *t*_imp_(*τ*). As expected the resulting *τ* * is reduced as we increase the temperature, reflecting the faster nature of the dynamics.

### Lorenz system

Our approach to state space reconstruction and extraction of coarse-grained dynamics is equally applicable to deterministic systems, which we demonstrate using the Lorenz equations,

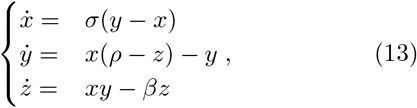

in the standard chaotic regime *σ* = 10, *ρ* = 28 and *β* = 8*/*3, Fig. 3(A). We take measurements of only the *x* variable (sampled at *δt* = 0.01 s for 10^5^ s), reconstruct the state space, and extract coarse-grained dynamics through the transfer operator. To reconstruct a maximally-predictive state space, we add delays to the measurement series to minimize the entropy rate, shown in Fig. 3(B) for different numbers of delays *K* and partitions *N*. We use *K*\* = 12 frames = 0.12 s, an embedding window for which the entropy rate is constant, Fig. S5(B), to reconstruct the state space. Unlike the stochastic dynamics within the double-well, the entropy rate of this deterministic dynamical system reaches an asymptotic value with increasing *N*, corresponding to the Kolmorgorov-Sinai entropy *h*_*KS*_ [62] (see also [39] for subtleties). Our result is in excellent agreement with the maximum Lyapunov exponent, as expected from Pesin’s theorem [63]. In addition to the entropy rate, our approach can easily convey other important invariants, such as the information dimension, Fig. S4.

**FIG. 3.**
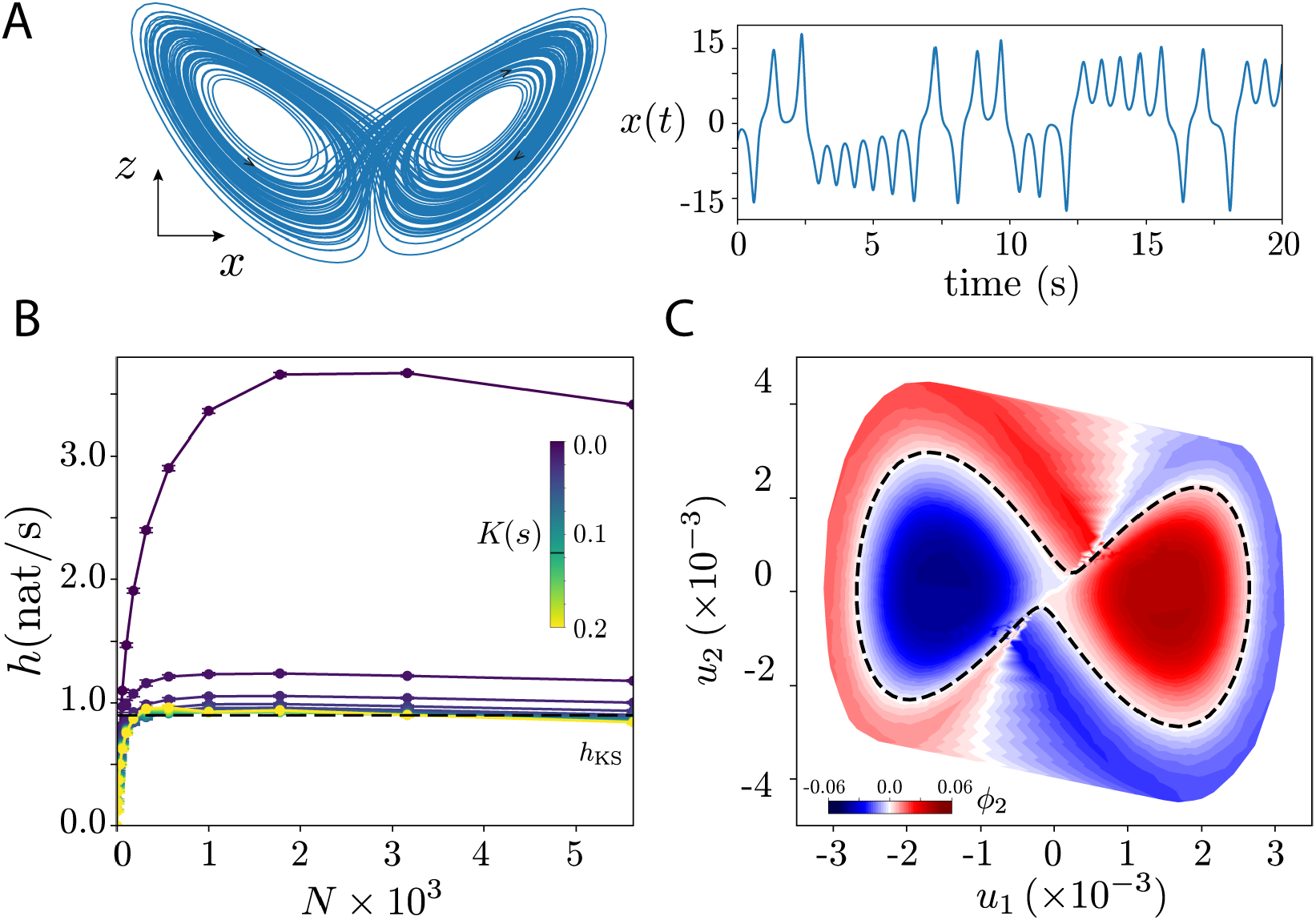
Maximally predictive ensemble dynamics in the Lorenz system. (A) We simulate the Lorenz equations, Eq. (13), in the standard chaotic regime and extract measurements of *x*(*t*), sampled at *δt* = 0.01 s for 5 × 10^5^ s. An example 100 s {*x, y*} trajectory is shown at left, and a sample 20 s measurement of *x* is shown at right. (B) The entropy rate *h* as a function of the number of partitions *N* for increasing number of delays *K* (color coded). From *K* ∼ 10*δt* = 0.1 s the entropy rate is approximately constant and we use *K*\* = 0.12 s for subsequent analysis. The estimated entropy rates converges to the maximum Lyapunov exponent *h*_*KS*_ = ∑_*λ>*0_ *λ* (dashed line, *h* = 0.91 (nat/s) [123]). (C) Contour plot of the first non-trivial transfer eigenvector of the inferred reversibilized transfer operator, *ϕ*_2_, projected along the largest two SVD modes of the reconstructed state space (*u*_1_, *u*_2_), computed with transition time *τ* * = 0.1 s and *N* = 10^3.5^ partitions. The sign of *ϕ*_2_ divides the state space into its almost invariant sets, which are partially split along the shortest unstable periodic orbit [64], here identified directly from the data using recurrences (Methods). Error estimates on the slowest relaxation time are 95% confidence intervals on the estimates obtained from 250 non-overlapping trajectory segments of duration 2000 s.

We further explore the connection to ergodic properties by showing a contour plot of the inferred first non-trivial eigenfunction *ϕ*_2_ of the reversibilized transfer operator *P*_*r*_, Eq. (9), constructed with *N* = 10^3.5^ partitions and transition time *τ* * = 0.1 s, and shown in the space of first two largest singular modes (*u*_1_, *u*_2_), Fig. 3(C). The sign of *ϕ*_2_ divides the state space into its almost invariant sets, Fig. S3(B), which are partially split along the shortest unstable periodic orbit (UPO) [64, 65], identified here through recurrences [66], Fig. S3(C) (Methods). Chaotic dynamics generally exhibit an intricate interplay of multiple timescales, which is apparent in the eigenvalue structure of *P*_*r*_ for different transition times, Fig. S3(A), which is more complicated than that of the double-well. Indeed, in the chaotic regime, the Lorenz dynamics can be described through a skeleton of UPOs [67], which are reflected in the periodic behavior of the eigenvalues as a function of *τ*.

### Identifying “run-and-pirouette” navigation directly from posture dynamics in *C. elegans*

Foraging behavior in the nematode worm *C. elegans*, an important model organism in genetics and neuroscience [68, 69], provides an excellent example of multiple timescales in biology; we apply our ensemble approach to demonstrate a direct bridge between short-time posture dynamics and the longer-lived states underlying navigation. On a 2D agar plate, worms move by making dorsoventral bends along their bodies [70], which are captured through a multidimensional time series of posture changes [71]. At the shortest timescales, posture dynamics are spatially organized as traveling waves along the body which give rise to forward, backward and turning locomotion [71–73]. Longer-lasting behaviors are performed by stringing together such body waves, and in navigation these have been classified phenomenologically into two elementary states: a series of forward body waves capable of lasting several seconds called “runs”, interrupted by sharp “pirouettes”, which are composed of backward and turning body waves [74]. We employ a previously analysed dataset [75] composed of 35 minute recordings of *N* = 12 N2 lab-strain worms freely moving on an agar plate, sampled at *δt* = 1*/*16 s. We measure the worm’s body posture using a low dimensional representation of the centerline, expressed as five “eigen-worm” coefficients 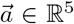 [71, 75], Fig. 4(A). To reconstruct the state space and transfer operators, we stack *K* delays of the measurement time series, and estimate the entropy rate as a function of *K* and the number of partitions *N*, Fig. 4(B). After *N* ∼ 1000 partitions we start observing finite-size effects as the entropy rate stops increasing, specially for lower *K*. Nonetheless, after *K* ∼ 8 frames = 0.5 s the entropy rate curves start collapsing, and we choose *K*\* = 11 frames = 0.6875 s to define the reconstructed space *X*_*K*\*_ as this corresponds to a regime in which the entropy rate is roughly constant, Fig. S5(C). We choose *N* = 1000 partitions so as to maximize the entropy rate and extract the *K* → ∞ limit of the entropy rate by extrapolation [76]: *h*_∞_ = 1.19 ± (1.11, 1.25) nat/s, obtained by extracting the offset of a linear fit in the interval 1*/K* = [1, 8] s^−1^. The early *K* behavior *h* is proportional to 1*/K*, indicating that the system has finite range correlations [43, 76], Fig. 4(B, inset). The large *K* decrease of *h* results from sampling error in the high-dimensional space [76]. Our estimated entropy rate is in good agreement with a previous calculation using estimated Lyapunov exponents from the same data [73]. Given *K*\*, we approximate the ensemble dynamics with *N* = 1000 partitions, and estimate the relaxation times of the first ten eigenfunctions of the reversibilized transition matrix, Fig. 4(C). We choose *τ* * = 0.75 s (dashed line) for subsequent analysis, the earliest *τ* where an approximately-Markov regime appears in the long-lived dynamics.

**FIG. 4.**
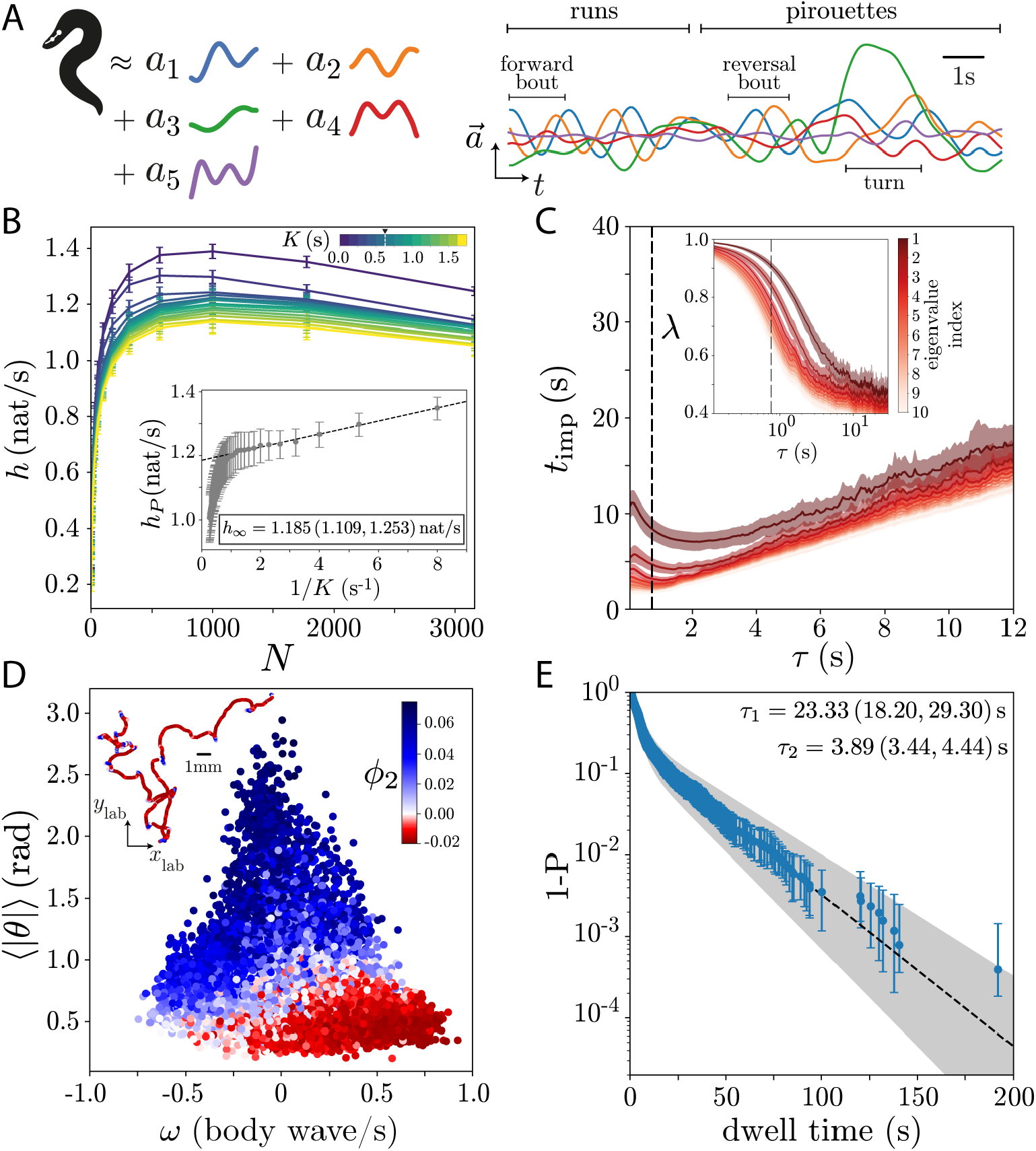
Deriving “run-and-pirouette” navigation from posture dynamics in *C. elegans*. (A-left) We represent the centerline posture dynamics of the worm’s body by a 5D (eigenworm) time series [71]. In food-free conditions, the short time scale behavior roughly consists of forward bouts, reversals and sharp turns. On longer time scales, worms (like other animals) combine basic behaviors into longer term strategies such as navigation, here exemplified by relatively straight “runs” interrupted by abrupt “pirouettes” [74]. (B) Entropy rate as a function of the number of delays *K* and partitions *N*. The curves collapse after *K* ∼ 0.5 s, indicating that the entropy rate is approximately constant. We choose *K*\* = 11 frames = 0.6875 s for subsequent analysis. For *N* ≳ 1000 partitions finite-size effects are apparent, and we extract the *K* → ∞ limit of the entropy rate by extrapolation *h*_∞_ = 1.19 ± (1.11, 1.25) nat/s (inset). Error bars are 95% confidence intervals of the mean, bootstrapped across worms. (C) The ten largest implied time scales as a function of transition time *τ*. We choose *τ* * = 0.75 s (dashed line) for subsequent analysis, 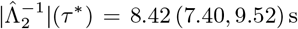. Error bars are 95% confidence intervals bootstrapped across worms. (inset) Eigenvalues of the transfer operator as a function of *τ*. As with previous examples, at very small *τ* or very large *τ* the eigenvalues collapse. We observe the largest separation for intermediate *τ*. (D) We use projections along *ϕ*_2_ to color the body wave phase velocities *ω* and mean body curvature ⟨|*θ* |⟩, averaged in 2*τ* = 1.5 s windows. Positive values align with negative phase velocities and large turning amplitudes, indicative of “pirouettes”, while negative values correspond to positive phase velocities and low turning amplitudes, indicative of “forward runs”. (inset) An example 10 minute long centroid trajectory color coded by *ϕ*_2_. Negative projections occur during “runs”, while positive values are found when worms abruptly change the locomotion direction. (E) We cluster *ϕ*_2_ into two coarse-grained states and find that the cumulative distribution of resulting run lengths 1 − *P* (*t*_state_ *< t*) is characterized by two time scales, which we extract by fitting a sum of exponential functions, and in agreement with previous phenomenological observations [74]. The timescale errors and error bars are 95% confidence intervals bootstrapped across events, while the shaded area represents the 95% confidence intervals bootstrapped across worms. The deviation at longer dwell-times foreshadows a finer subdivision, which we discuss in the context of Fig. 6.

To identify the longer-time behavior apparent in the transfer operator, we project the dynamics along *ϕ*_2_ (the 2nd eigenvector of the reversibilized operator) and use this mode to color the body wave phase velocities *ω* and mean body curvature ⟨| *θ* |⟩, averaged in 2*τ* * = 1.5 s windows, Fig. 4(D). Positive values (blue) generally align with negative phase velocities and large turning amplitudes, indicative of “pirouettes”, while negative values (red) correspond to positive phase velocities and low turning amplitudes, indicative of “forward runs”. In the inset we show an example 10 minute long centroid trajectory color coded by the projection along *ϕ*_2_. Negative projections occur during “runs”, while positive values are found when worms abruptly change the direction of locomotion. We cluster *ϕ*_2_ into two coarse-grained states by maximizing the coherence measure *χ*, Eq. (10), and find that the cumulative distribution of resulting run lengths 1 − *P* (*t*_state_ < *t*) is roughly characterized by two time scales, Fig. 4(E), fit by a sum of exponential functions and in excellent agreement with previous phenomenological observations [74].

The “run” and “pirouettes” state transition dynamics are out of equilibrium. The transition rates imply a relaxation time of 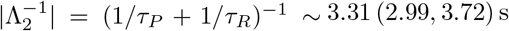, which is much shorter than the implied time scale of 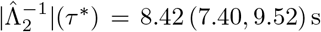, computed from the reversibilized dynamics. Indeed, positive entropy production has been reported in the short-time posture dynamics [73]. For out-of-equilibrium systems, we generally expect a discrepancy in eigenvalues (timescales) between *P* and *P*_*r*_, an aspect to which we return in Discussion.

### Operator-based simulations predict both fine-scale posture movements and large-scale trajectory diffusion

The inferred transfer operators provide a means for principled coarse-graining and a simple yet accurate model of the underlying dynamics. In the worm’s posture movement, we use *P*_*ij*_ to *simulate* symbolic sequences and show that these simulations capture fine-scale posture auto-correlation functions and large-scale state kinetics and diffusion of foraging trajectories.

We start from the initial discrete state of an individual worm and simulate symbolic sequences by sampling from the conditional probability distribution *P* (*s*_*j*_ |*ŝ*_*i*_), Fig. 5(A), where *ŝ*_*i*_ is the current state, *s*_*j*_ are all possible future states after a time scale *τ* * and *P* (*s*_*j*_|*ŝ*_*i*_) is the *i*-th row of the transfer operator matrix inferred for that worm *P*_*ij*_(*τ* *). The result is a sequence of microstates with the same duration as the worm trajectories, but with a sampling time *δt* = *τ* *. These dynamics are effectively diffusive in the state space: hopping between microstates according to the Markov dynamics and random selection from the population of state space points *X*_*i*_ within each visited partition.

**FIG. 5.**
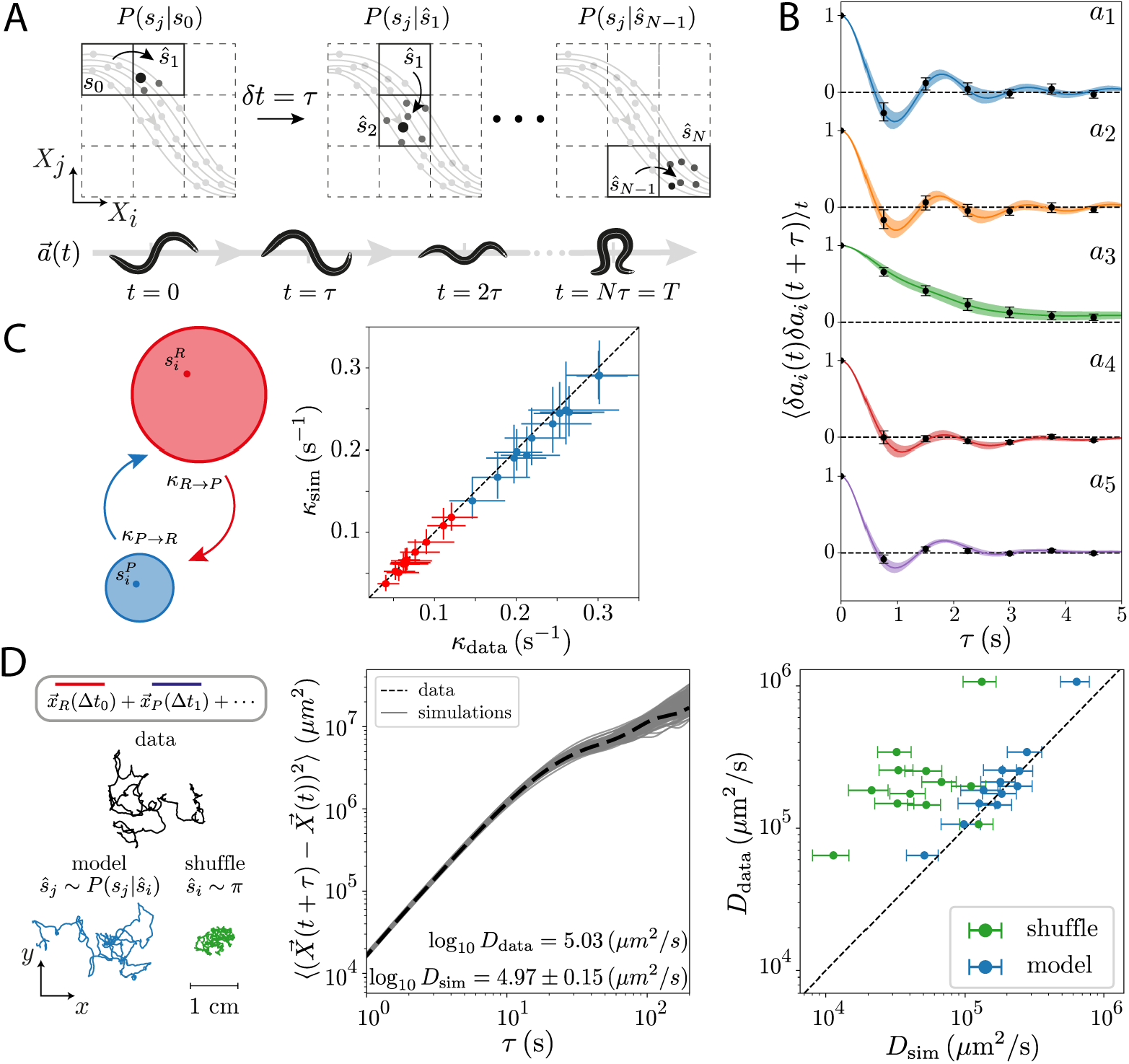
Operator-derived simulations predict both fine-scale posture dynamics and large-scale diffusion of foraging trajectories. (A) Schematic of the simulation method. Starting form the initial microstate of each worm *s*_0_, the next microstate is obtained by sampling from *P* (*s*_*j*_|*s*_0_) constructed with *τ* = *τ* * = 0.75 s. We add new microstates in the same fashion, resulting in a symbolic sequence of length *N* = *T δt/τ* *. Within each microstate, we randomly choose a point in the associated reconstructed state space, *X*, and use this point do define the body posture 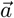. Through this effective diffusion in state space we obtain time series of postures through a sequence of “eigenworm” coefficients 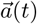 sampled at *δt* = *τ* *.(B) Simulated posture dynamics accurately captures the autocorrelation function of the “eigenworm” coefficient time series. Colors denote the different modes, while black dots correspond to the simulations. Error bars and shaded regions correspond to 95% confidence intervals on the estimate of the mean autocorrelation function bootstrapped across worms. (C) Simulated sequences exhibit kinetic rates between “run” and “pirourette” states in good agreement with data. We coarse-grain the symbolic sequence into run and pirouette states, as in Fig. 4, and show the mean transition rates from the run to the pirouette state *κ*_*R*→*P*_ (red) and from the pirouette to the run state *κ*_*P* →*R*_ (blue) obtained for each of the 12 recorded worms by inverting the mean dwell times in each state. Data error bars are 95% confidence intervals of the mean bootstrapped across run and pirouette segments, simulation error bars are 95% confidence intervals of the mean bootstrapped across 1000 simulations. (D) Centroid trajectories built from simulated symbol series exhibit diffusion constants and a ballistic-to-diffusive transition in agreement with data. (D-left) We simulate centroid trajectories by iteratively drawing from the space of actual trajectories, choosing a centroid run trajectory *X*_*R*_ (red) or a centroid pirouette trajectory *X*_*P*_ (blue) with duration Δ*t* which matches the dwell time of the microstate operator dynamics. We then append each new centroid segment to the end of the previous segment and align the direction of the worm’s movement across the boundary. We show the trajectory of an example worm, as well as simulated trajectories generated from the operator dynamics *ŝ*_*j*_ ∼ *P* (*s*_*j*_ | *ŝ*_*i*_) (blue), and from a shuffle which obeys the same steady-state distribution *ŝ*_*i*_∼ *π* (green). (D-middle) Mean square displacement of centroid trajectories obtained from 1000 simulations (gray) and the data (black dashed line) for an example worm undergoing a ballistic-to-diffusive transition. Errors of the diffusion coefficient are standard deviations across simulations. (D-right) Estimated effective diffusion coefficients obtained from simulations. We estimate *D* by fitting the slope of the mean square displacement curves in the interval *τ* ∈ [60, 100] s. Sequences that preserve the steady-state distribution but are otherwise random (green) are substantially different. Errors are standard deviations of the estimated diffusion coefficients across simulations.

From the diffusion in state space, where each state corresponds to a sequence of postures, we obtain a time series of “eigenworm” coefficients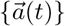 by sampling the first posture in each state: we thus obtain a simulated time series of postures of the same duration as the experimental data, sampled at *δt* = *τ* *. The autocorrelation functions of the simulated time series, Fig. 5(B), and the simulated mode distributions, Fig. S6, exhibits excellent agreement with worm data. On larger scales, we assign each microstate to a “run” or “pirouette” state as before, and estimate the average kinetic transition rates from runs-to-pirouettes *κ*_*R*→*P*_ and from pirouettes-to-runs *κ*_*P* →*R*_, finding close agreement between data and simulations, Fig. 5(C).

The accuracy of the Markov dynamics suggests that it is also possible to recover the diffusive properties of foraging trajectories. At the coarse-grained level, the symbolic sequences become binary sequences of run and pirouette states, with differing durations Δ*t*. We *generate* centroid trajectories by first sampling from the set of run and pirouette trajectories for each duration Δ*t*, Fig. S7(A). We then transform symbolic sequences into centroid trajectories by aligning and appending run and pirouette trajectories, sampled from the space of centroid trajectories with the appropriate duration (Methods). This yields simulated centroid trajectories that maintain the trajectory characteristics within each macrostate, but otherwise destroy correlations between them. In Fig. 5(D-left), we show an example centroid trajectory from the data (black), an example trajectory generated from the Markov model (blue) and an example trajectory generated from a random shuffle that obeys the stationary distribution *π* (green): the model generated trajectory is qualitatively more realistic than that of the shuffle, even though the shuffle respects the underlying fraction of runs and pirouettes.

To quantify the similarity between the simulated centroid trajectories and the data, we estimate the mean squared displacement 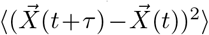 (MSD), which exhibits a transition between super-diffusive (nearly ballistic) and diffusive behavior between 10 s and 100 s [77– 80], Fig. 5(D-middle), Fig. S7(B). The operator-based simulations accurately capture the MSD across a wide range of scales, including the ballistic-to-diffusive transition. To further assess the quality of the simulations, we estimate an effective diffusion coefficient by fitting MSD = *Dτ* in the linear regime ^3^ and find that, across worms, the resulting diffusion coefficients obtained from simulations closely follow the data, Fig. 5(D-right). In contrast, centroid trajectories obtained from symbolic sequences that respect the underlying invariant density *π* but are otherwise random fail to mimic the extent to which worms explore the environment Fig. 5(D-left), which results in an underestimation of the effective diffusion coefficients, Fig. 5(D-right). Sampling from the underlying centroid trajectories is equivalent to having an accurate mechanical model of the interaction between the body wave and the medium and our results demonstrate that it is possible to go from microscopic posture dynamics to diffusive properties in a living organism. Despite fine-scale differences in the dynamics of each worm, a single *τ* * allows for an accurate prediction of the diffusive properties across the population.

### An operator-based “top-down” subdivision reveals additional states

While dividing the dynamics along *ϕ*_2_ identifies the longest lived states, splitting the free energy landscape along its highest energy barrier, what if there are important macroscopic states within each metastable set? We leverage the inferred transfer operator to perform a sequential subdivision of the state space, revealing finer-scale states, Fig. 6(A) and (Methods). At each step, the metastable state with the largest measure is subdivided along the first non-trivial eigenvector of the reversibilized transition matrix conditioned solely on the microstates within the metastable state. As before, we partition according to the corresponding first non-trivial eigen-function, maximizing the coherence measure *χ*, Eq. (10), defined only on the partitions of the highest measure macrostate. This yields a subdivision of the state space that obeys the structure of the free energy landscape; at each iteration, we subdivide the system along the largest energy barrier within the highest measure basin. Similar ideas have been used previously [82], where the notion of relatively coherent sets has been introduced. We note that our metastate subdivisioning process proceeds *from the longest-lived states down* rather than from the shortest-lived states up, where the latter is more common in temporal clustering approaches [83–85].

**FIG. 6.**
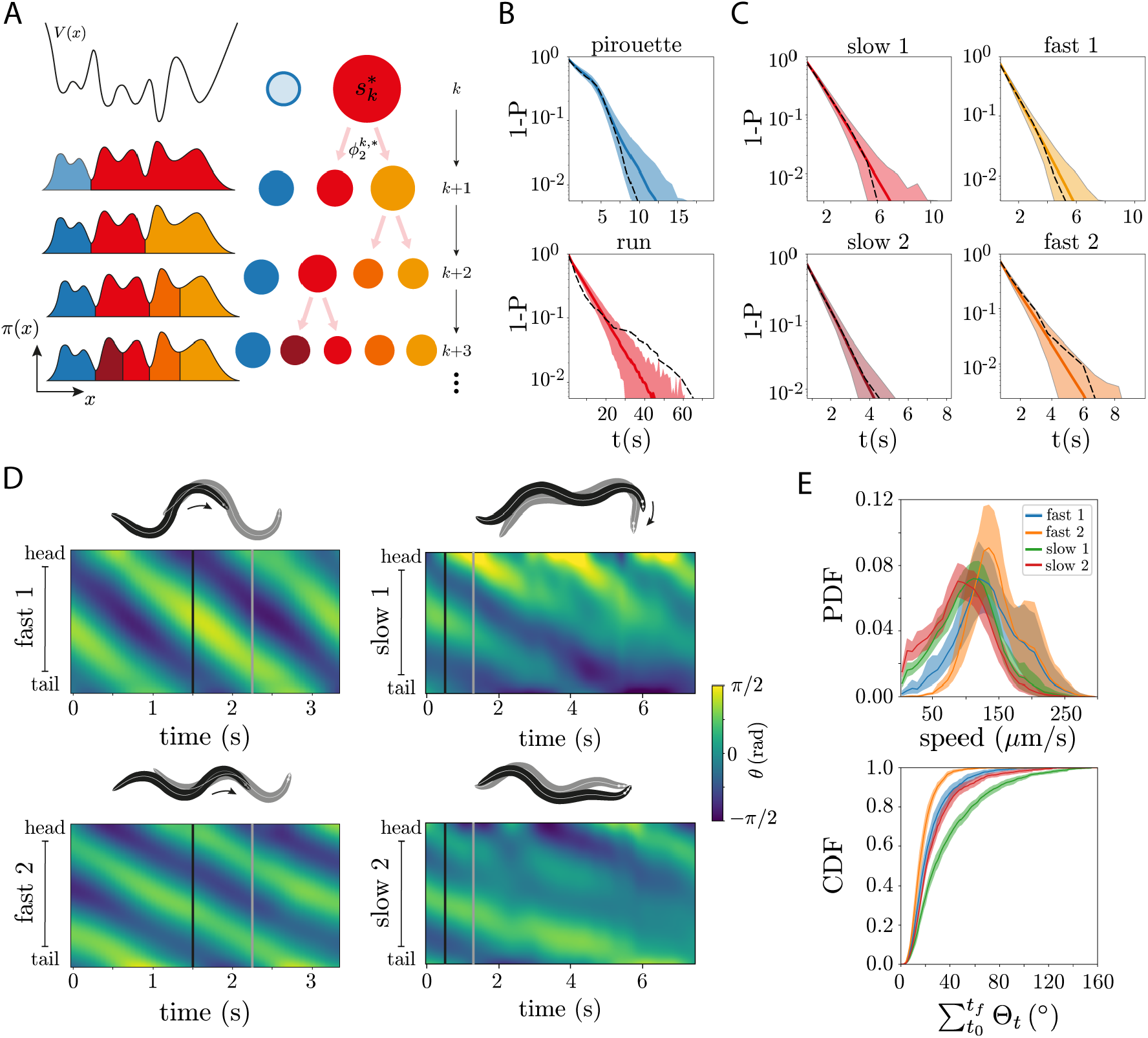
Operator-based subdivisioning reveals additional behavioral states. (A) State space subdivision. At each iteration *k*, we subdivide the highest measure metastable state, 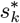 : we ignore the rest of the state space (shaded blue region) and partition along the first non-trivial eigenfunction obtained from the transition probability matrix conditioned only on the states within this metastable set, 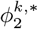 [82] (Methods). In principle, subdivisioning can be repeated as long as transitions between microstates are observed. (B) Dwell time distributions of the first two metastable states for simulations of the operator dynamics (blue - pirouette, red - run) and the data (black dashed line) for an example worm. The run state exhibits a heavy tail not accurately captured in our simulations. Shaded errors are 95% confidence intervals across 1000 simulations. (C) Dwell time distributions in the 4-state subdivision of the run state, for an example worm. Subdividing the run state into slower and faster forward movements better captures the dwell time distributions, see also Fig. S8(A). Shaded errors are 95% confidence intervals across 1000 simulations. (D-left) The two fast states correspond to distinct gaits. We show example body curvature *θ* kymographs with example postures taken *τ* = 0.75 s apart. (D-right) Example kymographs for the two slow states: “slow 1” exhibits head-casts [124], while “slow 2” has only minor movements, effectively a “pause”. (E) - Centroid-derived characterization of the four run states. (E-top) - Probability Distribution Function (PDF) of the centroid speeds for the four run states. Differences between fast states (blue and orange) and between slow states (green and red) are subtle from a centroid perspective, with the speed distributions nearly overlapping. (E-bottom) - Cumulative distribution function (CDF) of the summed angle between subsequent centroid velocity vectors measuring the overall curvature of the path. The higher wavelength observed in “fast 1” states results in a gradual “weathervaning” [125, 126] of the trajectory when compared to the trajectories of the “fast 2” state which are comparably straighter. Similarly, the “head-casts” in the “slow 1” state result in the gradual slow reorientation of the worm, when compared to the “slow 2” state for which trajectories are less curved. Shaded areas correspond to 95% confidence intervals bootstrapped across worms. To avoid spurious large angles due to reversals, we discard the 10% largest magnitude angles in all states.

We illustrate this approach with an application to the worms’ posture dynamics, where finer-scale states can be particularly relevant in connection with the underlying genetic and neural control machinery. In fact, while the operator derived dynamics accurately capture the average kinetic rates in the 2-state coarse-graining, the dwell time distribution of run states exhibits, in most worm recordings, heavy tails not captured by a 2-state subdivision, as exemplified in Fig. 6(B) and summarized in Fig. S8(A). To explore the possibility that these heavy tails are indicative of a finer decomposition relevant for worm behavior, we systematically subdivide the metastable states,

Beyond the initial subdivision into runs and pirouettes, the next iterations yield 4 distinct run states: first the run state is split into fast and slow runs, and then each of these is subdivided into two distinct states. This 5-state subdivision allows us to better capture the dwell time distributions Fig. 6(C), with a Kolmogorov-Sinai test across worms showing that the dwell time distributions obtained from simulations are indistinguishable from the data with a 5-state partition, whereas with a 2-state partition some dwell time distributions in the run state are significantly different from the data, Fig. S8(A). The four “run” states reveal important fine scale differences in the body wave dynamics, Fig. 6(D, E), further detailed in Fig. S8(B, C). The subdivision of the fast states yields distinct gaits with distinct body wavelengths but comparable speeds; we show the body curvature as function of time for two example runs of the fast state in Fig. 6(D-left) and the respective speeds Fig. 6(E-top). The longer wavelength state typically results in higher curvature runs compared to the other fast state, Fig. 6(E-bottom). Accordingly, the largest wavelength fast state exhibits larger coiling amplitudes when compared to the second fast state. The slow states exhibit larger amplitude head and tail excitations (as measured by the fourth eigenworm coefficient, *a*_4_), when compared to the fast states. In addition, the first slow state exhibits a dorsal bias (negative *a*_3_), which results in the gradual reorientation of the worm through head-casts, Fig. 6(D-right). In contrast, the second slow state is more akin to a pure pause state, for which there are only minor posture changes that do not result in a significant displacement of the worm, Fig. 6(D-right). Such fine-grained states are essential in capturing the posture kinematics, but have not yet been characterized in the *C. elegans* literature, and add an important component for further quantitative understanding in this important and accessible model organism.

## DISCUSSION

When seeking order in complex systems, many simplifications are possible, and this choice has consequences for both the nature and complexity of the resulting model. Often such simplifications are chosen *a priori*, and the complexity of the model is taken to be a fact of the underlying dynamics. But this is not generally true; the definition of the states and the resulting dynamics between them are inextricably linked. Here we choose the simplification informed by the resulting model dynamics; we define states in an attempt to achieve a Markov model of the state dynamics. While this class may seem restrictive, we find models of the dynamics that are accurate across a range of time scales and systems. Even for dynamics that do not ultimately permit a Markovian description, we offer a principled coarse graining to independently investigate long time scales.

Applicable to both deterministic and stochastic systems, we combine state space reconstruction with ensemble dynamics to infer probabilistic state space transitions directly from data, which we apply to the Langevin dynamics of a particle in a double-well, the Lorenz system, and the wiggling of the nematode worm *C. elegans*. Rather than seek low-dimensonal descriptions of the data directly, as is often the choice in cluster-based analysis (e.g. [79, 86–88]), we instead first *expand* in representation complexity: enlarging the putative state to short sequences and constructing a *maximum* entropy partition to include as much predictive information as possible in the embedding. For dynamical systems with a continuous state space, maximizing the partition entropy can also approximate a generating partition, which preserves ergodic characteristics [89].

By characterizing nonlinear dynamics through transitions between state space partitions, we trade trajectory-based techniques for the analysis of linear operators. This approach is complementary to previous methods of state space reconstruction which focus on geometric and topological quantities such as Lyapunov exponents, the Kolmogorov-Sinai entropy and dimensions [33, 66, 73, 90, 91]. In particular, trajectory-based techniques of ergodic analysis rely on precise estimates of dimension and of the local Jacobian, which can be challenging in systems with unknown equations.

In our approach, we recognize and address the mutual dependence of state space reconstruction and Markovian evolution: reconstructing the state space is required for an effective Markovian description and the framework of ensemble dynamics provides the principled, information-theoretic measure of memory used to optimize the reconstructed state. Previous work in molecular and neural dynamics [29, 92] have also implicitly expanded states in time in order to improve the accuracy of Markov models. By reconstructing the state space, hidden dynamics are included into the state, such that the slow modes uncovered by the operator reconstruction correspond to the slowest possible ergodic dynamics in the system. However, slow non-ergodic degrees of freedom can render the data non-stationary, in which case the transfer operator has an explicit time dependence 𝒫 _*τ*_ (*t*). Nonetheless, a *τ* -parametrized family of coherent sets can capture *moving* regions of the state space that remain coherent within a time scale *τ* [52, 93–97].

To optimally identify almost invariant sets, we use the reversibilized operator *P*_*r*_, Eq. (9), which is generally different from *P*. For an overdamped system in thermodynamic equilibrium, this symmetrization simply enforces detailed balance and 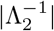 is directly related to the hopping rates between metastable states. Irreversible dynamics, however, result in complex eigenvalues, which cannot be directly associated with relaxation time scales. Generally, the dynamics of *P*_*r*_ will relax more slowly than those of *P*, providing an upper bound to the true relaxation times [98, 99]. We thus assume that the Markovianity of the metastable dynamics of *P* is reflected in *P*_*r*_, and we find that is a good approximation in our examples, Fig. S9. It will also be interesting to explore approaches based directly on *P* [52].

When the state-space is known or its reconstruction decoupled from the ensemble dynamics, mesh-free discretizations have been used to characterize ensemble evolution, including diffusion maps [92, 100] and other kernel based approaches, such as Reproducing Kernel Hilbert Spaces [101] or Extended Dynamic Mode Decomposition [102]. Though powerful, such methods require subtle choices in kernels, neighborhood length scales and transition times, for which we lack guiding principles.

A particularly interesting focus of maximally-predictive ensemble dynamics is in the quantitative understanding of animal behavior [15, 16, 103]. This may seem paradoxical; organism-scale behavior arises as the cumulative output from a wide range of influences, including genetic, neural, and bio-mechanical systems, all of which may in principle introduce important hidden-states (see e.g. [104, 105]) and thus non-Markovian dynamics [87, 106]. Yet, the process of constructing behavioral states has often been decoupled from the analysis of the dynamics and “under-embedding” could easily introduce apparent memory.

In the behavior of the nematode *C. elegans*, maximally predictive ensemble dynamics connects sub-second posture dynamics to long-lasting, compound behavioral states. The worm embedding combines postures across roughly a quarter of the duration of a worm’s typical body wave and results in delay and entropy estimates in agreement with previous work [73]. On longer timescales, the posture-based “run-and-pirouette” navigation strategy [74, 107] derived from the operator dynamics provide an accurate and principled discretization of foraging behavior, disentangling motions that are confounded by centroid-derived measurements (see e.g. [108]). For example, the different “gaits” that result from the subdivision of the run state exhibit comparable centroid speeds and are only clearly distinguished due to the different body wavelengths. These emergent Markov dynamics offer a promising and powerful demonstration of quantitative connections across the hierarchy of movement behavior generally exhibited by all organisms [87, 109]. Particularly interesting future directions include the analysis of even longer dynamics in *C. elegans* [110–113].

The principled integration of fluctuating, microscopic dynamics resulting in coarse-grained but effective theories is a remarkable success of statistical physics. Conceptually similar, our work here is designed towards systems sampled from data whose fundamental dynamics are unknown. We leverage, rather than ignore, small-scale variability to subsume nonlinear dynamics into linear, ensemble evolution, enabling the principled identification of coarse-grained, long-timescale processes, which we expect to be informative in a wide variety of systems.

## ACKNOWLEDGEMENTS

We thank Massimo Vergassola and Federica Ferretti for comments. This work was supported by OIST Graduate University (TA, GJS), a program grant from the Netherlands Organization for Scientific Research (AC, GJS), by the Herchel Smith Fund (DJ), and by Vrije Universiteit Amsterdam (AC, GJS).

## METHODS

### Software and data availability

Code for reproducing our results is publicly available: https://github.com/ AntonioCCosta/predictive_ensemble_dynamics/. Data can be found in [114].

### State space reconstruction

Given a measurement time series, 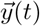, with *t* ∈ {*δt*, …, *Tδt*} and 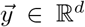, we build a trajectory matrix by stacking *K* time-shifted copies of 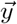, yielding a (*T* − *K*) × *Kd* matrix *X*_*K*_. For each *K*, we partition the candidate state space and estimate the entropy rate of the associated Markov chain (see below). We choose *K*\* such that 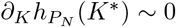.

### State space partitioning

We partition the state space into *N* Voronoi cells, *A*_*i*_, *i* {1, …, *N*}, through k-means clustering with a k-means++ initialization using scikit-learn [115].

### Approximation of the Perron-Frobenius operator

We build a finite dimensional approximation of the Perron-Frobenius operator using an Ulam-Galerkin discretization. A Galerkin projection takes the infinite dimensional operator onto an *N*×*N* operator of finite rank by truncating an infinite dimensional set of basis functions at *N*. Ulam’s method uses characteristic functions as the basis for this projection,

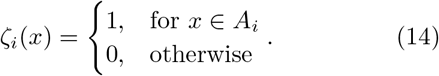

Our characteristic functions are implicitly defined through the k-means discretization of the space. We thus partition the space into *N* connected sets with nonempty and disjoint interior that covers 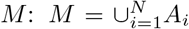, and approximate the operator as a Markov chain by counting transitions from *A*_*i*_ to *A*_*j*_ in a finite time *τ*. Given T observations, a set of *N* partitions, and a transition time *τ*, we compute

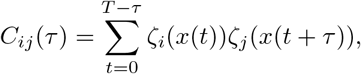

where *ζ*_*i*_(*x*) are the Ulam basis functions, Eq. (14). The maximum likelihood estimator of the transition matrix is obtained by simply row normalizing the count matrix,

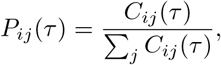

which yields an approximation of the Perron-Frobenius operator.

### Invariant density estimation

Given a transition matrix *P*, the invariant density is obtained through the left eigenvector of the non-degenerate eigenvalue 1 of *P, πP* = *π*: *π*_*i*_ is the probability of finding the system in a partition *A*_*i*_.

### Entropy estimation

Given a partition of the state space into regions of size *ϵ*, the probability of the system visiting *M* consecutive cells is given by the joint probability 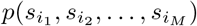, where *i*_*k*_ ∈ {0, …, *N*} and *N* is the number of partitions of the state space. The *M* –block Shannon entropies are defined as the sum over possible sequences, 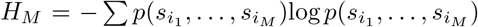, measured in *nats*. As in a thermodynamic system, for which entropy grows with the system size, the entropy of a symbolic sequence grows with the sequence length. The rate of entropy growth is defined through the entropy of the condition probability of the system visiting the cell *s*_*M*+1_ given that it has visited the previous *M* cells: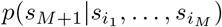,

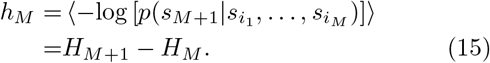

The limit *h* = lim_*M*→∞_ *h*_*M*_ is the entropy rate of the source and measures the overall unpredictability of the dynamics. It is a non-decreasing function of number of partitions *N*, being bounded from above by *h* ≤ log*N*. To characterize the underlying dynamical process, one takes the supremum of the entropy rate over all possible partitions, thus obtaining the Kolmogorov-Sinai entropy rate,

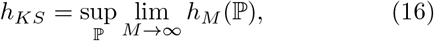

where ℙ is a candidate partition of the state space.

One approach to the estimation of the entropy rate, more common among the information theory community, is to estimate the *M* -block entropy rates for increasing *M*, given a fixed small number of partitions *N*. This entails counting all sequences of length *M* and estimating the joint probabilities 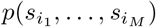. However, high dimensional non-linear dynamical systems generally require a large number of partitions to approach the Kolmogorov-Sinai limit. Given this constraint, computing sequence probabilities for reasonably large sequence lengths *M* quickly becomes infeasible since the number of possible sequences grows with *N*^*M*^.

Another approach, more common in the dynamical systems literature, is to build blocks of *K* delays and estimate correlation entropies, introduced by Cohen and Procaccia (CP) [38]. Given a measurement time series {*y*(*t*)} where *t* ∈ {*τ*, 2*τ*, …, *Tτ*}, *ϵ*-tubes are built around sequences of length *Kτ, y*^*K*^(*t*), and the entropy is estimated as,

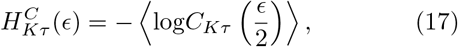

Where 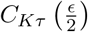 is the correlation function of *K*-dimensional *ϵ*-tubes built from time series measurements with a sampling time of *τ*, which essentially measures the probability of finding two *y*^*K*^(*t*) vectors within a distance *ϵ* of each other. This approach avoids the counting problem of direct estimates for large *N* (small *ϵ*) and *K*. The correlation entropy rate can then be obtained by estimating,

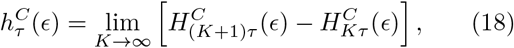

which converges to the Kolmogorov-Sinai entropy in the limit *ϵ* → 0 and *τ* → 0. It is important the note that correlations are typically measured using a Chebyshev distance, 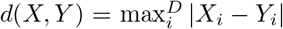 for *X,Y* ∈ ℝ^*D*^ and | · | the absolute value, and so the precise values of 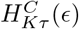 may depend on the chosen metric.

As described in the main text, we present here a hybrid approach to the estimation of entropy. As in the CP method, we add time into the definition of state (instead of increasing the length of the sequence of discrete states), but then partition that state space and use the inferred Markov chain to approach the entropy rate of the source for increasing K, Eq. (4).

We illustrate the similarities between our partition based approach and the geometric approach using an autoregressive process of order *p*, AR(p), as in [116]. The behavior of the entropy as a function of the number of delays is an indication of the memory of the autoregressive process. We use a simple AR(2) model, with

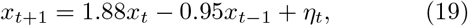

where *η*_*t*_ is a delta correlated Gaussian random variable with unit variance. In Fig. S10 we show the resulting estimates of the entropy rate (top) and predictive information (bottom) using the CP method of Eq. (18) [38] and our partition based approach Eq. (4). The behavior of the entropy as a function of the number of delays is qualitatively similar between the two methods, despite the dramatic differences in metrics and estimation procedures. The entropy rate grows faster with the decreasing length scale for *K* = 1 and the behavior of *h* with the length scale for *K* ≳ 2 is *h*(*K, ϵ*) = *h*(∞,*ϵ*) log *ϵ* with identical *h*(∞, *ϵ*). The difference between *h*(*K* = 1, *ϵ*) and *h*(*K* = 2, *ϵ*) stems from the memory of the process. Similar conclusions can be taken by looking at the predictive information estimates Fig. S10(bottom), *I*_pred_ = *H − h* (Eq. 6): the system exhibits a higher degree of predictability when *K* > 1, indicating that it is a second order process. We can therefore recover the dimensionality of the underlying state space, which in this case simply corresponds to the order of the autoregressive process.

### Identifying metastable states

Metastable states correspond to regions of the state space that the system visits often, separated by regions where transitions occur. We therefore search for collections of states that move coherently with the flow: their relative future states belong to the same macroscopic set. As discussed in the Introduction, we leverage a heuristic introduced in [47], which makes use of the first non-trivial eigenfunction of the transfer operator to identify almost invariant sets. As in [64], we will work with a time reversibilized transfer operator 𝒫_*r*_: the coherence properties are invariant to this transformation [56] and the analysis is simplified due to the optimality properties of reversible Markov chains. In discrete time and space, *P*_*r*_ is defined as

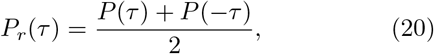

where

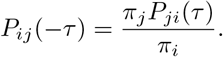

is the stochastic matrix governing the time-reversal of the Markov chain. The first nontrivial (*λ* < 1) right eigenvector of *P*_*r*_, *ϕ*_2_, allows us to define the macrostates as

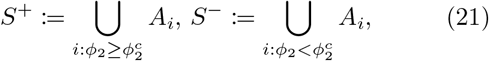

where 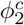 is chosen so as to maximize *χ*, Eq. (10), and 𝒳_*μ,τ*_ are computed in a discretized state space,

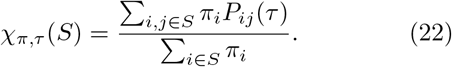

In practice, we compute Eq. (10) for all 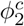 (*S*^+^ and *S*^−^ are implicit functions of 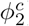) and find the global maximum: this is an inexpensive calculations since 𝒳 can be obtained by matrix multiplications and 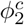 can only take *N* different values, where *N* is the number of partitions. We note that for an overdamped thermodynamic system in equilibrium, the eigenvectors *ϕ*_*k*_ are discrete approximations of the eigenfunctions of the Koopman operator [102], which are nearly constant within a metastable set thus simplifying the clustering into almost invariant sets.

### Operator-based state space subdivision

We leverage the notion of relatively coherent sets [82] to subdivide the state space. However, instead of subdividing both metastable state at each iteration, we identify the state with the most measure *S*′ and build a new transition matrix only with partitions belonging to that state,

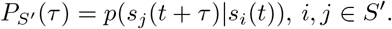

From *P*_*S*′_ we proceed as before: we compute the stationary distribution of *S*^′^ through the first left eigenvector of *P*_*S*′_, 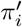, build the corresponding reversibilized transition matrix *P*_*r,S*′_ and identify relatively metastable states through its first non-trivial eigenvector by maximizing Eq. (8) where *π*_*i*_ and *P*_*ij*_(*τ*) are replaced by their relative counterparts 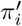 and *P*_*S*′_. The iterative process is illustrated in Fig. 6(A).

### Choice of transition time*τ* *

We choose *τ* * as the shortest transition time scale after which the inferred implied relaxation times reach a plateau. For *τ* too short, the approximation of the operator will yield a transition matrix that is nearly identity (due to the finite size of the partitions and too short transition time), which results in degenerate eigenvalues close to *λ* ∼ 1: an artifact of the discretization and not reflective of the underlying dynamics. For *τ* too large, the transition probabilities become indistinguishable from noisy estimates of invariant density, which results in a single surviving eigen-value *λ*_1_ = 1 while the remaining eigenvalues converge to a noise floor resulting from a finite sampling of the invariant density. Between such regimes, we find a region with the largest time scale separation (as illustrated in Fig. 1(C-middle)) which also corresponds to the regime for which the longest relaxation times *t*_imp_, Eq. (7), are robust to the choice of *τ*. We compute *t*_imp_ using the eigenvalues of the reversibilized transition matrix, *P*_*r*_, which only gives an upper bound to the relaxation dynamics. In addition, we assume that *P*_*r*_ can only be Markovian if also *P* is Markovian, a statement that holds for the systems studied here, Fig. S9.

### Matrix diagonalization

The high dimensionality and the sparsity of the transition matrices for large *N* results in numerical errors when using a naive estimator for the full spectrum of eigenvalues. In addition, since we are interested in the longest lived dynamics, we focus on finding only the *n*_modes_ largest magnitude real eigenvalues using the ARPACK [117] algorithm.

### Periodic Orbit identification

We identify the shortest period unstable periodic orbit of the Lorenz system, Eq. (13), by studying the distribution of recurrence times. We set a short distance *ϵ* and look for the times *p* at which ‖*X*_*t*+*p*_ − *X*_*t*_‖ < *ϵ*, where ‖. ‖ represents the Euclidean distance. In practice, we compute 1*/* ‖*X*_*t*+*p*_ − *X*_*t*_‖^2^ and find peaks with height larger than 1*/ϵ* ^2^ using the `find_peaks` function from the scipy.signal package [118]. For short enough *ϵ* (*ϵ* = 5 × 10^−6^ in our case), the distribution of recurrence times has its first peak at the period of the shortest unstable periodic orbit [119]. A trajectory corresponding to a shadow of this unstable periodic orbit is shown in Figs. 3C, S3C [120, 121].

### *C. elegans* centroid trajectory simulations

Given a symbolic sequence, we generate centroid trajectories by appending and aligning “run” and “pirouette” trajectories independently drawn from the set of all possible trajectories of a given duration. In the rare occasion that the generated symbolic sequences yield state durations that are longer than all the observed trajectories, we append multiple randomly sampled trajectories up to the corresponding duration. We align the beginning of a new sampled centroid trajectory with the end of the previous one in the following way: we average the velocity vectors in the 2*τ* * = 1.5 s segment preceding the sampled trajectory, estimate its orientation angle and rotate the trajectory such that the initial orientation matches the angle defined from the ending 2*τ* * segment of the previous trajectory. Trajectories with a duration less that 2*τ* * were discarded as they do no allow for an appropriate alignment. This process generates continuous centroid trajectories with the same overall duration as the experiments, but with independent segments randomly sampled from the population of “run” or “pirouette’ trajectories of a given duration.

### Relaxation times for the non-reversibilized dynamics

We choose *τ* * through the *real* spectrum of the reversibilized transition matrix *P*_*r*_, looking for a constant implied relaxation time scale 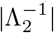, which sets an upper bound to the underlying relaxation dynamics. We thus make an indirect test for the Markovianity of *P*, assuming that *P*_*r*_ is only Markovian when *P* is also Markovian. Estimating time scales directly from *P* is non-trivial due to the possibility of complex eigenvalues. However, we can probe the relaxation dynamics by estimating the first passage time distribution between coarse-grained states directly from the full Markov chain. Coarse-graining along the first non-trivial eigenvector of the reversibilized transition matrix *ϕ*_2_, we obtain two macrostates *a* and *b* with measures *π*_*a*_ and *π*_*b*_ respectively. The dwell time distribution in these states *f*_*α*_(*t*) can be obtained directly from the full non-reversibilized Markov chain *P* : the system would have to hop from *β* to *α*, stay in *α* for *t* = *nτ* steps and then hop back to state *β*, which yields 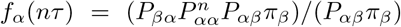, where *P*_*βα*_(*τ*) is the probability of transitioning from a microstate in macrostate *α* to a microstate in macrostate *β* in a transition time *τ, α, β* ∈ {*a, b*}. The mean first passage time can then be obtained by integrating 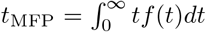 and the decay rate to the invariant density is given by |Λ_2_| = 1*/t*_*a*→*b*_ + 1*/t*_*b*→*a*_. However, this calculation is challenging to do numerically, due to the large number of matrix multiplications needed to reach the *t* = *nτ* → ∞ limit and the low accuracy of the Riemann integral for large *τ*. An equivalent yet simpler calculation can be done by leveraging the eigenvalue decomposition of the infinitesimal dynamics: the first passage time distribution simply becomes 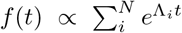, and the mean first passage time yields 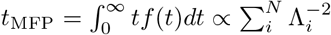 where Λ_*i*_ ∈ℝ^+^ are the eigenvalues of the infinitesimal generator of the dynamics. For a discrete spectrum with a time scale separation, the mean first passage time will be governed by the first non trivial eigenvalue, Λ_2_ ∼ 0; 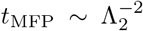, since the other eigenvalues Λ_*i*_ ≪ 0 for *i* > 2 only add minor contributions. Therefore, a simpler yet equivalent estimate of the slow kinetic rates can be obtained by first coarse-graining the system into its macrostates *a* and *b*, and then computing the first non-trivial eigenvalue of the coarse-grained two-state Markov chain *P*_*c*_, which will be a real number. The corresponding relaxation times for our example applications are shown in Fig. S9.

### Double-well simulation

We use an Euler-Maruyama integration scheme to simulate Eq. (12) and generate a *T* = 10^6^s long trajectory of a particle in a double-well potential with *m* and *γ* set to unity and *β*^−1^ = {0.5, 0.75, 1.00, 1.25, 1.5, 1.75, 2.00, 2.25, 2.5} J. We first sampled at 100Hz, and then downsampled to a sampling time of *δt* = 0.05 s. In addition, the first 1000 s were discarded to avoid transients. In S2(A), we project the Boltzmann distribution into the SVD space spanned by [*u*_1_, *u*_2_] by learning a linear mapping *θ*, between [*u*_1_, *u*_2_] and [*x, v*]: [*u*_1_, *u*_2_] = *θ* · [*x, v*]. This allows us to project the centroids of each Voronoi cell in the [*u*_1_, *u*_2_] space to the [*x, v*] space, and estimate the corresponding Boltzmann weight *p*([*u*_1_, *u*_2_] · *θ*^−1^) = *p*([*x, v*]) = exp{−*β*[(*x*^2^ − 1)^2^ + *v*^2^*/*2}*/Z*, where *Z* represent the normalization.

### Lorenz system simulation

We use scipy’s odeint package [118] to generate a *T* = 5 × 10^5^ s long trajectory of the Lorenz system in the standard chaotic regime (*σ, ρ, β*) = (10, 28, 8*/*3), sampled at *δt* = 10^−2^ s^−1^. We discard the first 10^3^ s to avoid transients.

### *C. elegans* foraging dataset

We used a previously-analyzed dataset [71], in which N2-strain *C. elegans* were imaged at *f* = 32 Hz with a video tracking microscope on a food-free plate and downsampled to *f* = 16 Hz to incorporate coiled postures [75]. Worms were grown at 20^°^*C* under standard conditions [122]. Before imaging, worms were removed from bacteria-strewn agar plates using a platinum worm pick, and rinsed from *E. coli* by letting them swim for 1 min in NGM buffer. They were then transferred to an assay plate (9 cm Petri dish) that contained a copper ring (5.1 cm inner diameter) pressed into the agar surface, preventing the worm from reaching the side of the plate. Recording started approximately 5 min after the transfer, and lasted for 2100 s, for a total of *T* = 33600 frames. In Fig. 4(B), we infer the asymptotic limit of the entropy rate *h*_∞_ by linear extrapolation of 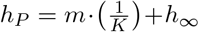, using the values of *h*_*P*_ obtained for each *K*, Fig. 4(B). The parameters *m* and *h*_∞_ were inferred by linear regression in the regime *K* ∈ [0.125, 1] s using the scikit-learn package in Python [115].

## SUPPLEMENTARY MATERIAL

**FIG. S1.**
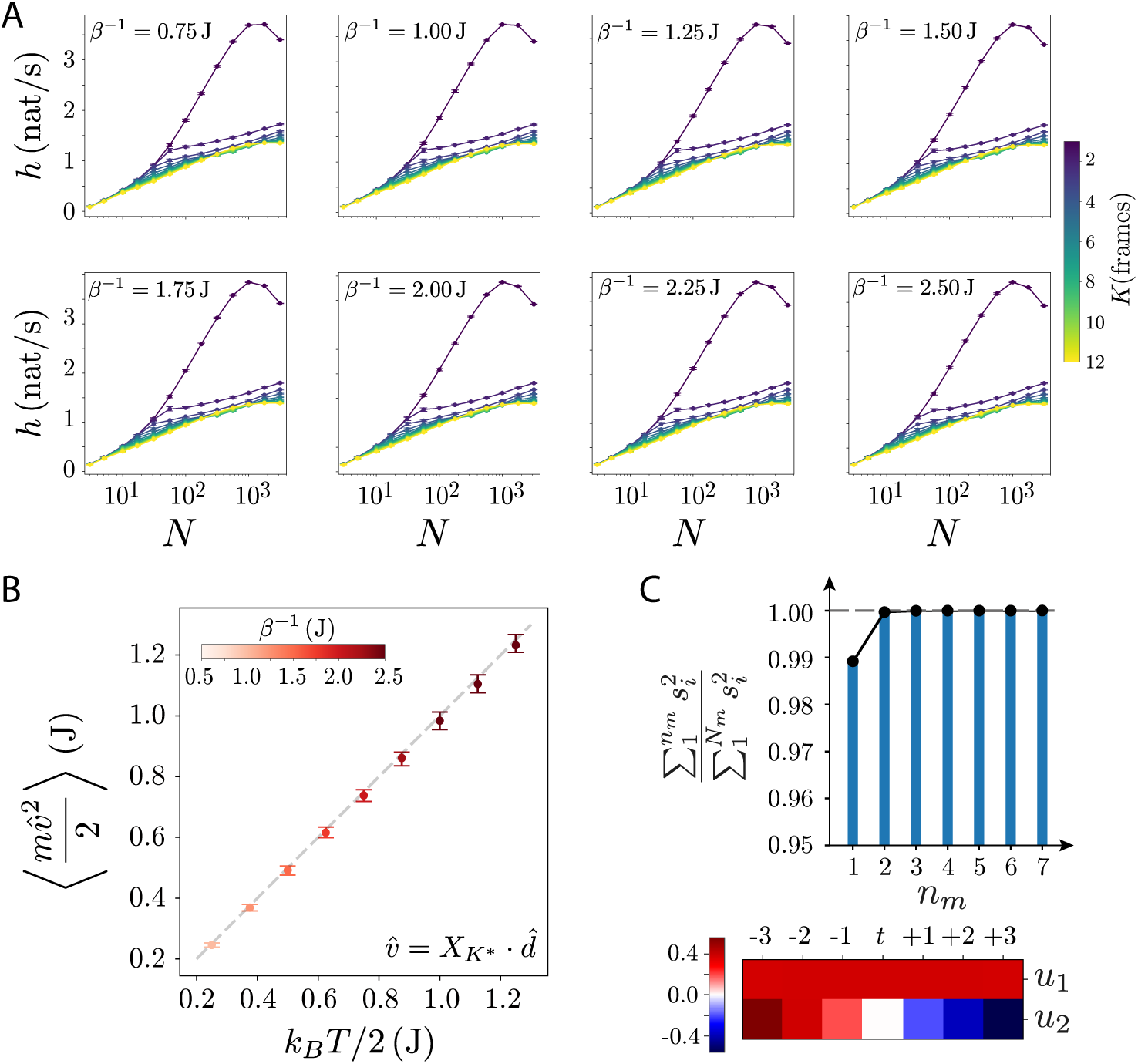
Details of the state space reconstruction of an underdamped particle in a double well potential. (A) Entropy rate as a function of delay *K* and number of partitions *N* across temperatures. The qualitative behavior of the entropy rate is notably conserved across temperatures. (B) We recover the equipartition theorem in the reconstructed state space, an indication that complete information is contained in the reconstructed state space. We use a 6th-order finite difference approximation to estimate the velocity 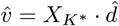 from the reconstructed state *X*_*K*_* with *K*\* = 7 frames, where 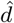 represent the 6th-order finite difference stencil. We then compute the average kinetic energy and show that it closely matches the expected value of (*k*_*B*_*T*)*/*2 from equipartition. We use *K*\* = 7 frames across temperatures, as the behavior of the entropy rate with *K* and *N* exhibits is similar across the range of temperatures studied here, Fig. S1(A), S5(A). Error bars show 95% bootstrapped confidence intervals of the mean velocity bootstrapped over 500 s time points. (C) Singular value decomposition of the reconstructed state space 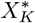 with *K*\* = 7 frames. The first two SVD modes capture nearly 100% of the variance (top). In addition, the resulting singular vectors capture the position and the velocity degrees of freedom (bottom): *u*_1_ is essentially a weighted sum of the position, while *u*_2_ is a derivative filter of the position, yielding an estimate of the local velocity.

**FIG. S2.**
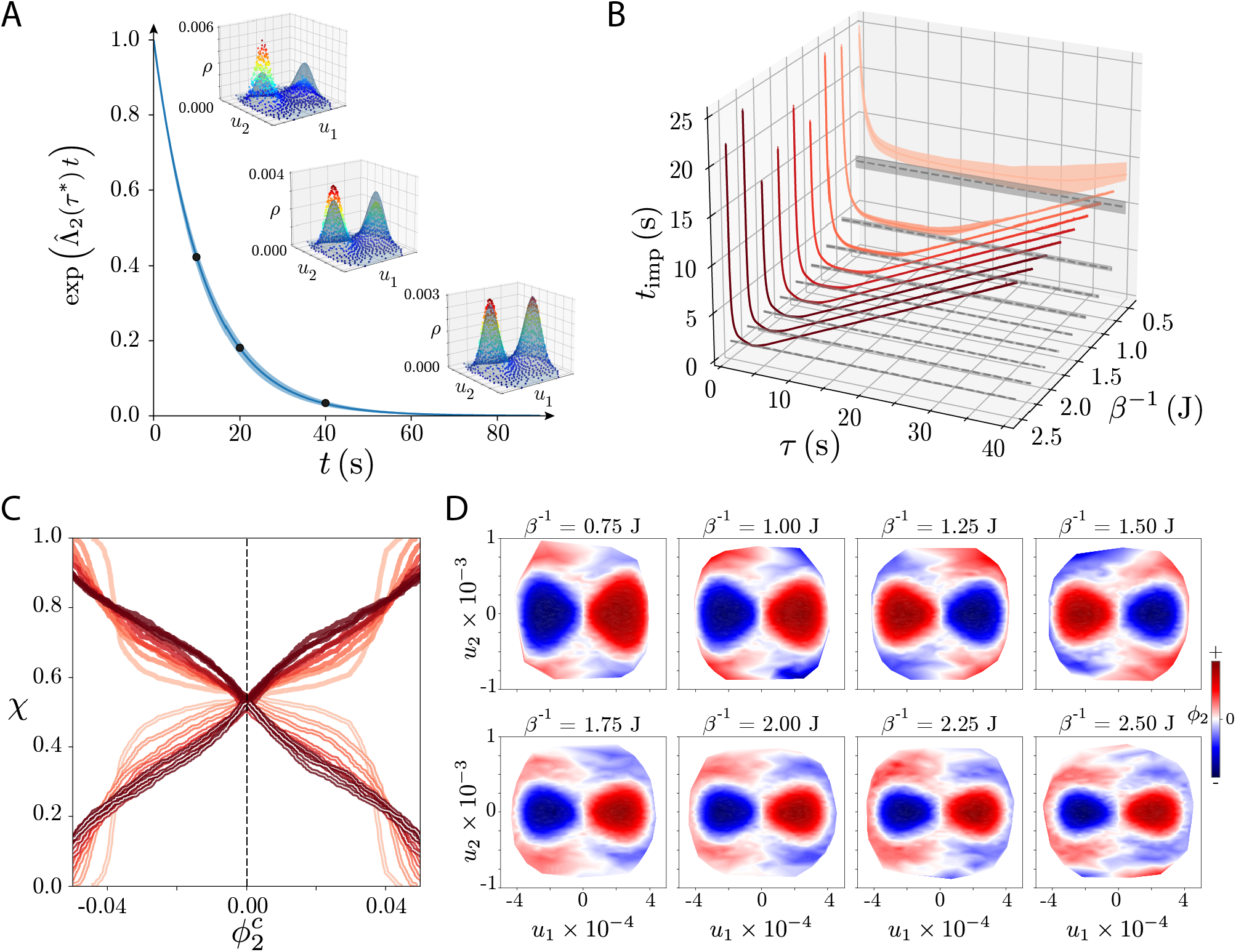
Transfer operator dynamics in the Double Well potential and metastable state identification across temperatures. (A) Ensemble dynamics captured by *ϕ*_2_ projected onto the largest two singular vectors (*u*_1_, *u*_2_) of the reconstructed state space for *β*^−1^ = 0.5 J, which match the position and velocity degrees of freedom, Fig. S1(C). The decay captures the slowest evolution towards the equilibrium density. We start from an initial ensemble sharply concentrated on a single partition on the bottom of the left well, and propagate the density using *P* with *β*^−1^ = 0.5 J. (B) Inferred longest implied time scales *t*_imp_ as a function of transition time *τ* and effective temperature *β*^−1^ = *k*_*B*_*T*. The qualitative behavior of *t*_imp_ with *τ* is conserved across temperatures: early transient decay to the time scale of the hopping rate *t*_MFP_*/*2 (gray dashed line) and long time linear increase with *τ* indicating independence of the states. Increasing the temperature has the expected effect of reducing the duration of the plateau: higher temperatures result in faster mixing, reducing the time it takes before states become independent and *t*_imp_ grows linearly. Accordingly, we observe a decrease in *t*_imp_ with temperature. We leverage the conserved qualitative behavior of the *t*_imp_ curves across temperatures to choose a consistent *τ* * as the time scale that minimizes *t*_imp_. This results in a lower *τ* * for higher temperatures, which reflects the faster nature of the dynamics: the fast modes relax faster, enabling for a larger time scale separation at shorter *τ*. (C) Coherence measure *χ*, as defined in Eq. (10) as a function of the inferred first non-trivial eigenfunction of 𝒫_*r*_, *ϕ*_2_, for different temperatures. For each temperature, we show both *χ*_*μ,τ*_ (*S*^+^) and *χ*_*μ,τ*_ (*S*^−^) as well as the minimum between them, *χ*, in a gray dashed line. Changing *τ* * to reflect the nature of the dynamics yields a relatively constant maximum *χ* across temperatures. Notably, at *τ* * the coherence measure indicates that nearly half the measure has escaped the metastable states. (D) *ϕ*_2_ projected onto the first two SVD modes of the reconstructed state *X*_*K*_ across temperatures: the slow eigenfunction mainly splits the state space along the position coordinate and the additional high velocity |*u*_2_| transition regions.

**FIG. S3.**
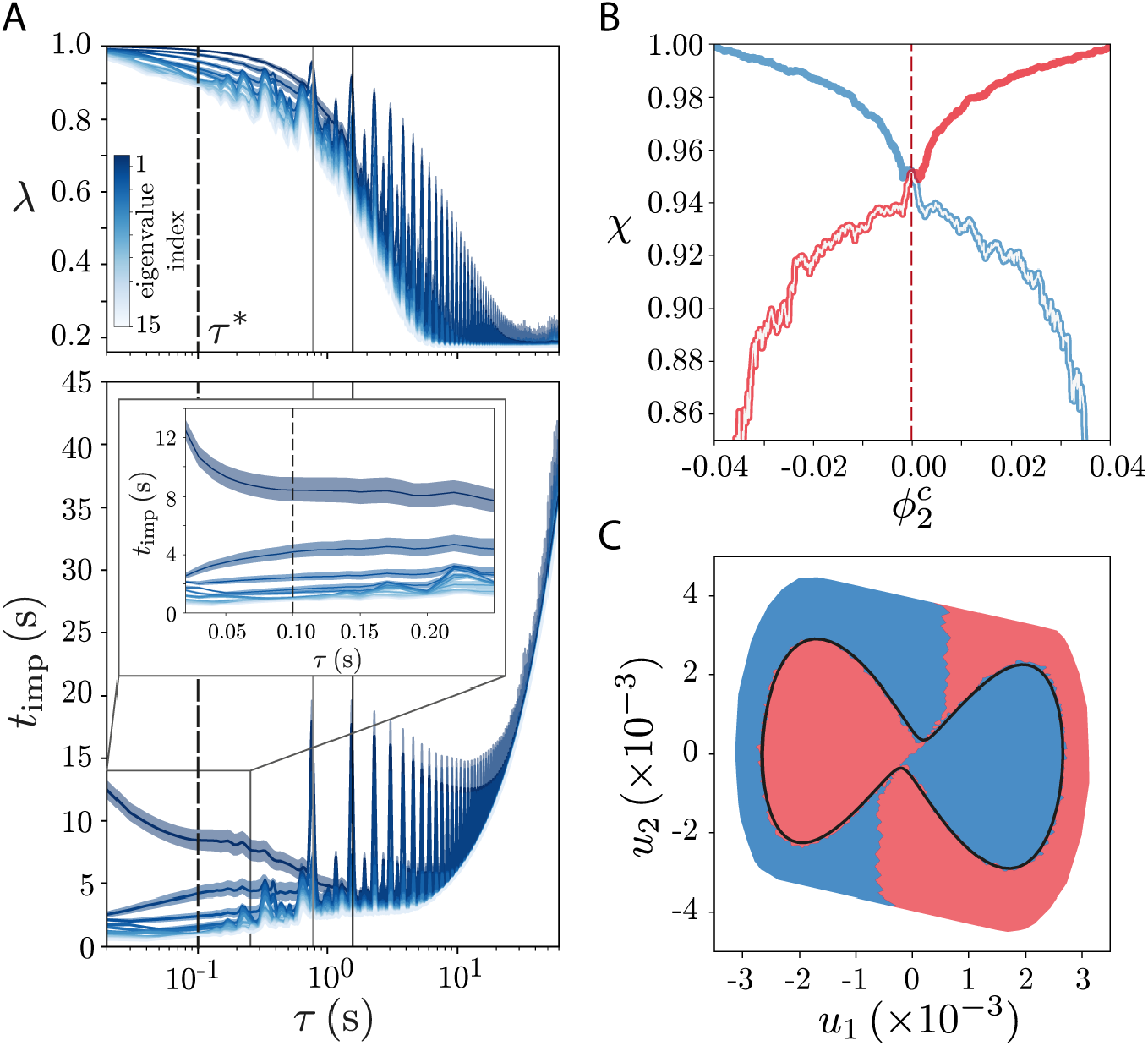
Details of the transfer operator analysis of the Lorenz system. (A) Behavior of the implied time scales *t*_imp_ with transition time *τ* for the Lorenz system. (A-top) 15 largest magnitude non-trivial eigenvalues as a function of *τ*. For very small *τ* the eigenvalues are nearly degenerate around 1. After this initial transient, the eigenvalues spread, but unlike the Double Well example the time scale separation is much less evident since we are in presence of deterministic chaos. For *τ* 0.75 ∼ s, we observe an increase in the inferred eigenvalues resulting from the periodicity of the Lorenz attractor: the black solid line represents the period of the shortest identified unstable periodic orbit (UPO) *T*_UPO_ = 1.55 s [119] (see Methods), while the gray solid line represents half of this period. The period of the UPO sets the onset of the increase in the estimated eigenvalues: the periodicity induces an aliasing effect; the transfer operator becomes again nearly identity and we observe the same behavior at integer multiples of *T*_UPO_*/*2. For very large *τ*, the transfer operator is composed of near copies of the stationary distribution, resulting in the collapse of the eigenvalues, as observed for the Double Well potential, Fig. 2(C-inset). (A-bottom) Implied relaxation times as a function of *τ*. As expected from the behavior of the eigenvalues (top), there is an initial short transient before the longest relaxation time settles into a near constant value (inset). For larger *τ*, the effects of the periodicity of the Lorenz attractor come into play: the time scale separation shortens and we observe peaks and integer multiples of *T*_UPO_*/*2. Despite the cyclic pattern, for large *τ* we observe a linear growth of the implied relaxation time scales as in the Double Well example (here seen as exponential growth on a logarithmic scale), Fig. 2(C). We choose *τ* * = 0.1 s yielding an upper bound on the mixing time of 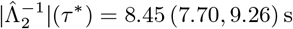. (B) Degree of almost invariance as measure by *χ*, Eq.(10). Colors represent the contribution of each almost invariant set (*χ*_*μ,τ*_ (*S*^+^) and *χ*_*μ,τ*_ (*S*^−^)) while the white line is the minimum between sets, *χ*. We split the state space along the global maximum of *χ* ∼ 0. (C) Resulting almost invariant sets, projected onto the first two singular vectors of the trajectory matrix, *X*_*K*_ and plot of the shortest UPO identified through recurrences (see Methods). The boundary between almost invariant sets is partially defined by the stable manifold of the UPO [64], of which we show only the orbit itself for clarity. Error bars represent 95% confidence intervals from 250 trajectories of duration 2000 s.

**FIG. S4.**
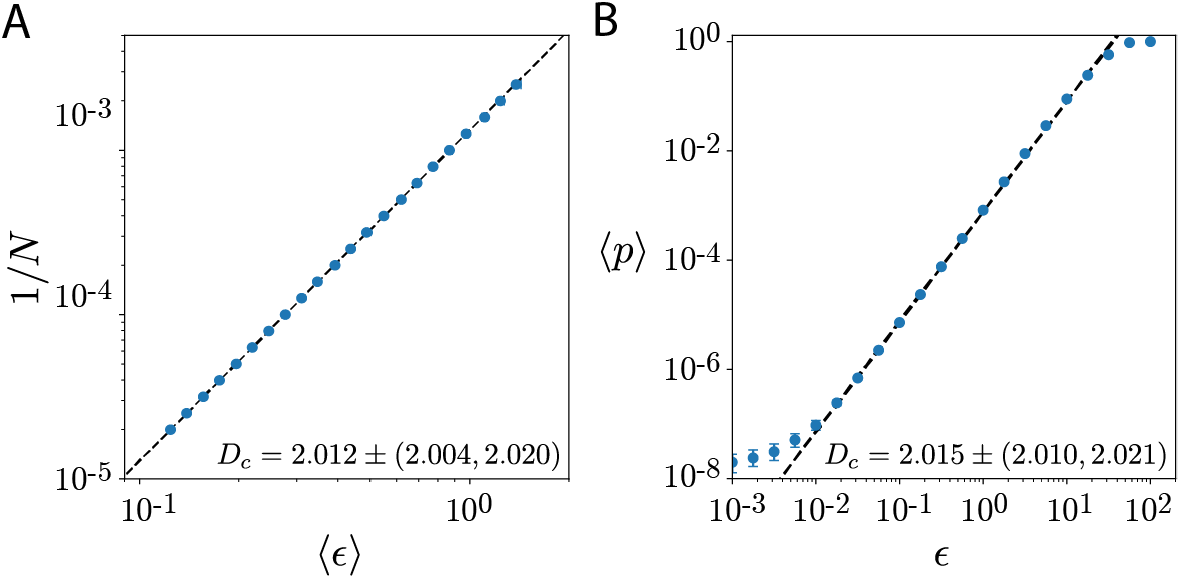
Estimation of the information dimension in a partitioned state space. The information dimension can be obtained by studying the scaling of the probability distribution with length scale. The average probabilities ⟨*p*⟩ of finding a neighbor within a length scale *ϵ* scales as 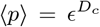, in which *D*_*c*_ is the correlation dimension (or pointwise dimension [40]), which generally bounds the information dimension from above *D*_*c*_ ≥ *D*_*I*_. For a typical chaotic attractor, and in the limit *ϵ* →0, the correlation dimension is indistinguishable from the information dimension, and thus 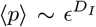. Estimates of information dimension are typically done using Chebyshev distances: 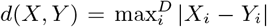 for *X, Y* ∈ ℝ^*D*^ and | · | the absolute value. (A) Information dimension estimates in a partitioned state space. The probability of belonging to a partition scales with the inverse of the number of partitions, and thus 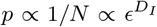. Therefore, we can directly estimate the information dimension by measuring how the within cluster distances scale with increasing *N*. We estimate *D*_*I*_ by linear regression of 1*/N* ∝ *D*_*I*_ log⟨ *ϵ* ⟩, where ⟨·⟩ represents the median. Error estimates for *D*_*I*_ are obtained by bootstrapping the median within-cluster distances. Error bars in ⟨ *ϵ*⟩ are generally smaller than the marker size. (B) Geometric estimation of the correlation dimension using the Cohen-Procaccia (CP) method [38] in our Lorenz system simulation. The CP method assesses the scaling of the probability distribution with length scale, by measuring the correlation function of a collection of state space points. We sample 5000 points from the attractor, and count the number of neighbors within a distance *ϵ* from the sampled points using a Chebyshev distance. *D*_*I*_ is then estimated by linear regression log⟨*p*⟩ ∝ *D*_*I*_ log *ϵ*. Error bars represent 95% confidence intervals of the estimate done 100 times with different 5000 point samples.

**FIG. S5.**
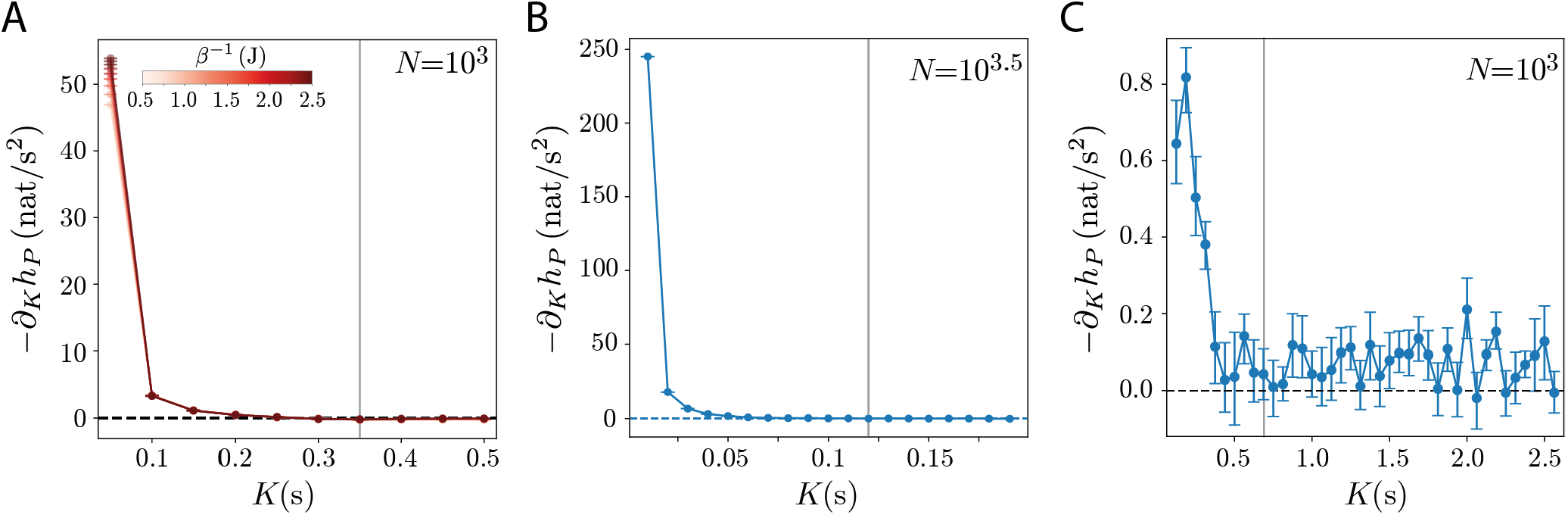
Change in entropy rates as a function of delays *K* for the double well example, the Lorenz system and *C. elegans* posture data. (A) Double well dynamics for *N* = 10^3^ partitions. The entropy rate reaches a plateau after *K* ≳ 6 frames = 0.3 s and we choose *K*\* = 7 frames = 0.35 s. Notably, the change in entropy rate is still significantly nonzero at *K* = 2 frames, reflecting the non-Markovian effects of the Euler integration scheme as discussed in the main text. Error bars represent 95% confidence intervals bootstrapped over 50000 s trajectory segments. (B) Lorenz system for *N* = 10^3.5^ partitions. The entropy rates reaches a plateau after *K* ≳ 0.05 s and we choose *K*\* = 0.12 s. Error bars represent 95% confidence intervals bootstrapped over 2000 s trajectory segments. (C) *C. elegans* posture dynamics for *N* = 10^3^ partitions. The entropy rates reaches a plateau after *K* ≳ 0.5 s and we choose *K*\* = 0.6875 s. Error bars represent 95% confidence intervals bootstrapped across worms.

**FIG. S6.**
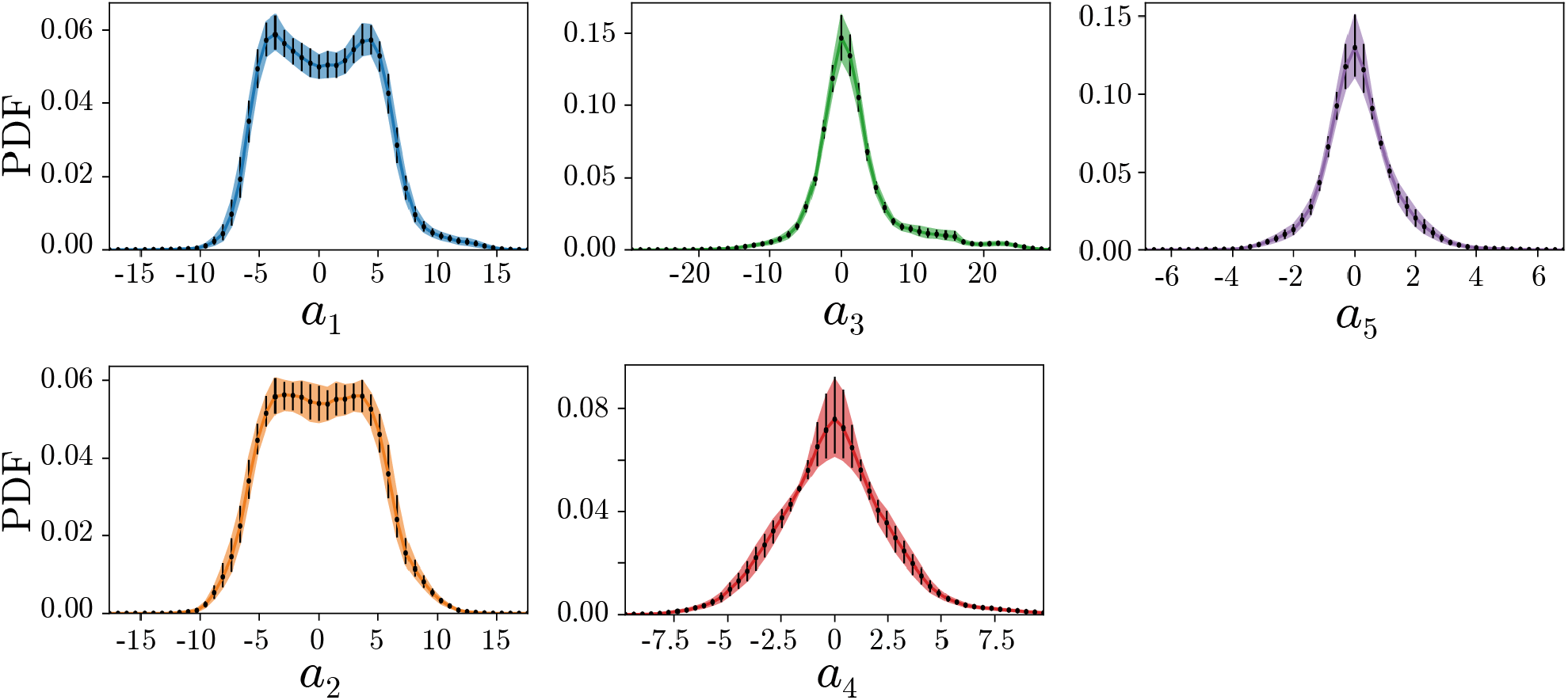
Probability Density Function (PDF) of the “eigenworm” coefficients show a tight agreement between the data (colors) and simulations (black error bars). Error bars correspond to 95% confidence intervals bootstrapped across worms.

**FIG. S7.**
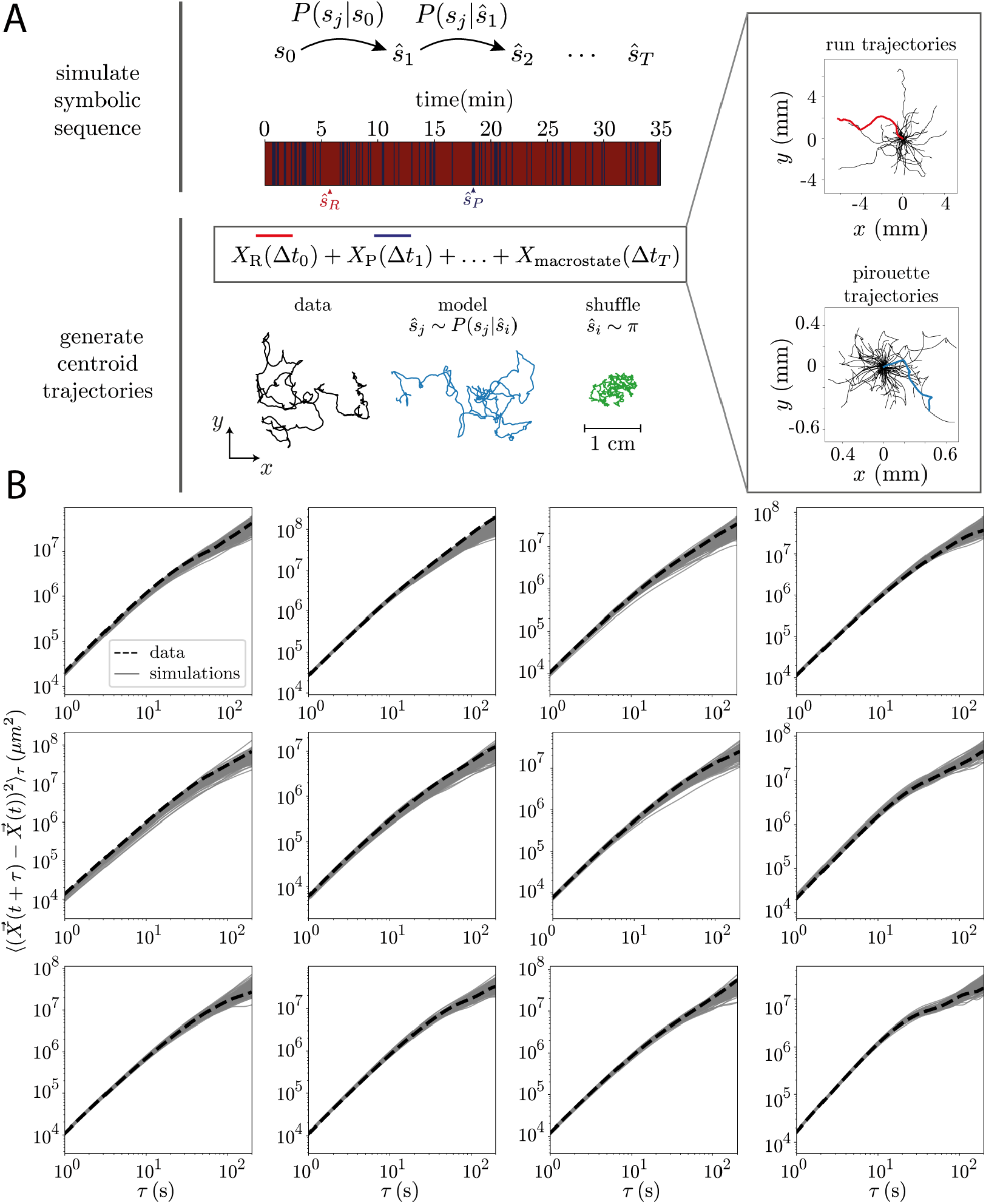
Details of the centroid trajectory simulations. (A) Illustration of the simulation method. (top) Starting from the initial condition *s*_0_, a partition on the reconstructed state space, the next state is sampled using the conditional probability *P* (*s*_*j*_ |*s*_0_). We iterate until we get a symbolic microstate sequence of the same length as the worm recordings. (middle) We associate each microstate with a “run” (red) or “pirouette” (blue) macrostate in which the microstate is contained, resulting in a binary sequence of macrostates with duration Δ*t*_*i*_. (right) We construct the simulated centroid trajectory by iteratively drawing from the space of actual trajectories, choosing a centroid run trajectory *X*_*R*_ (red) or a centroid pirouette trajectory *X*_*P*_ (blue) with duration which matches the dwell time resulting from the miscrostate operator dynamics. (bottom) We append each new centroid segment to the end of the previous segment and align the direction of the worm’s movement across the boundary. We thus retain the trajectory characteristics within each macrostate but destroy any correlations across them. We show the trajectory of an example worm, as well as simulated trajectories generated from the operator dynamics *ŝ*_*j*_ ∼ *P(s*_*j*_ | *ŝ*_*i*_) (blue), and from a shuffle which obeys the same steady-state distribution *ŝ*_*i*_ ∼ *π* (green). (B) We show the mean square displacements for the data (dashed line) of each worm, as well as for 100 centroid trajectory simulations generated from symbolic sequences simulated with the Markov model (gray). By fitting a linear function in the interval *τ* ∈ [60, 100] s we obtained the effective diffusivity estimate of Fig. 5(C-right).

**FIG. S8.**
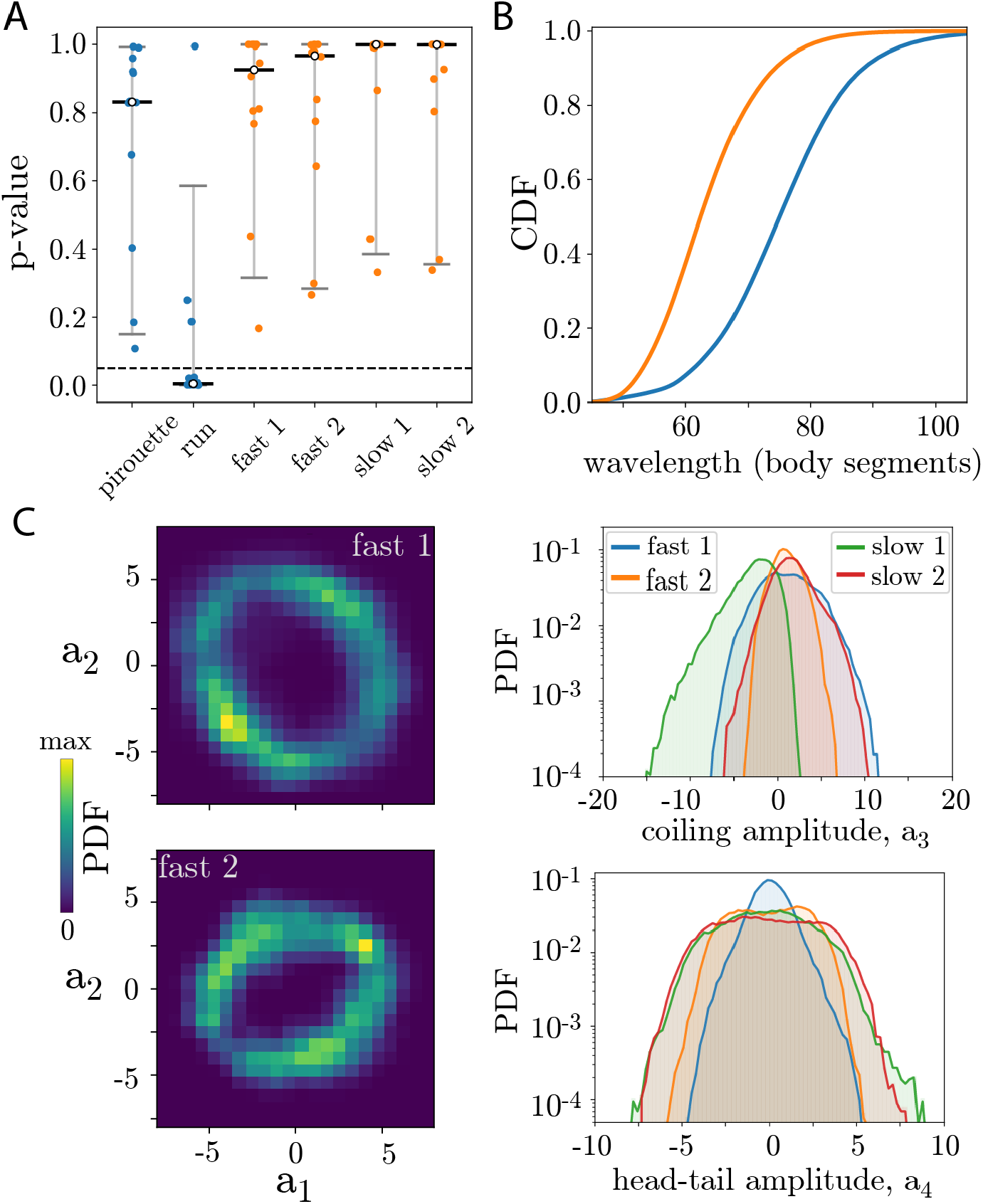
Details of the operator-guided subdivision of *C. elegans* postural dynamics. (A) Scatter plots of the *p*_values_ of a Kolmogorov-Smirnov test comparing the dwell time distributions obtained from simulations and the data for a 2-state (blue) and a 5-state (orange) coarse-graining, across worms. In the two state partition, a large fraction of the dwell time distributions in the run state are not accurately captured by the simulations (*p*_value_ *<* 0.05, dashed line), whereas subdividing the run state yields dwell time distributions that are statistically indistinguishable from the data. Vertical bars represent the median, 5-th and 95-th percentiles and we add horizontal Gaussian jitter to the scatter plots to aid visualization. (B) Cumulative Distribution Function (CDF) of the body wavelength in the two fast states, obtained as the peak in the spatial power spectrum (obtained using Welch’s method). The first fast state exhibits larger wavelength, spanning almost the entire body, whereas the second slow state exhibits shorter wavelengths. (C-left) The difference in wavelength in the two fast states is translated into different cyclic patterns in the (*a*_1_, *a*_2_) plane, which captures the worms’ body wave. Such subtle differences likely become more evident through the state space reconstruction. (C-right) Probability distribution of the coiling amplitude (*a*_3_, top) and head-tail amplitude (*a*_4_, bottom), across run states. The state “fast 1” exhibits a higher coiling amplitude and a lower head-tail amplitude, while “fast 2”, which displays straighter centroid paths, exhibits a lower coiling amplitude and higher modulation of the head-tail amplitude. The two slow states exhibit distinct biases in coiling amplitude: “slow 1” has a clear dorsal bias (*a*_3_ < 0) while “slow 2” has a slight ventral bias (*a*_3_ > 0). Both also exhibit more activity in the head-tail mode compared to the fast states.

**FIG. S9.**
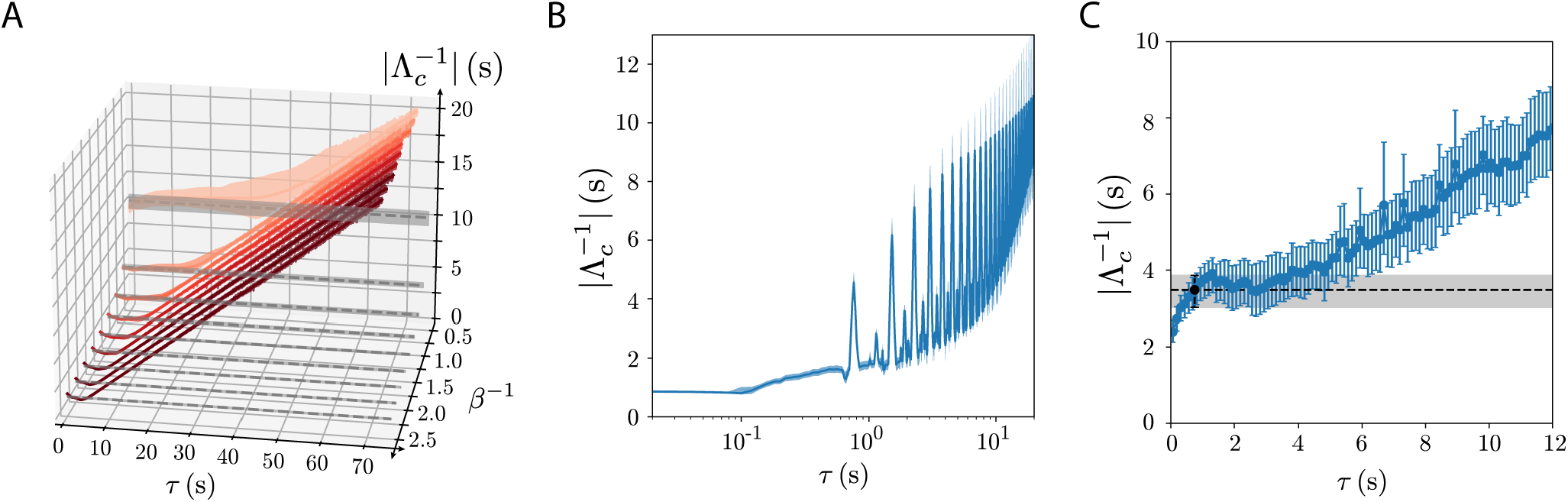
Markovianity of the irreversible Markov chain as measured through the inferred relaxation times. The reversibilized transition matrix provides an optimal partition into almost invariant sets, but the resulting kinetics does not necessarily capture the underlying dynamics. To directly probe the Markovianity of the underlying slow dynamics, we have to estimate the relaxation times for the non-reversibilized transition matrix, which should not change with *τ* when the dynamics is Markovian. However *P* generally presents complex eigenvalues and so the relaxation times are not directly attainable through spectral analysis. To solve this problem in systems which display a time-scale separation, we approximate the slow relaxation dynamics by using the metastable states to build a two-state, coarse-grained Markov chain *P*_*c*_, which necessarily has only real eigenvalues *λ*_*c*_ ∈ ℝ (details in Methods). The corresponding relaxation time is then obtained through 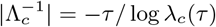. In general, we find that the regime in which 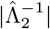 from *P*_*r*_ is constant overlaps with regime in which 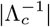 is also constant, although generally 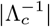 exhibits a larger Markovian regime than 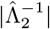. (A) Double Well example across temperatures, *β*^−1^. The time scales of *P* converge to the hopping time scale even at *τ* = *δt*, unlike 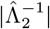 from *P*_*r*_. The underdamped nature of the dynamics creates microscopic fluxes that are erased by the reversibilization of Eq. (9), resulting in the overestimation of the relaxation times measured through *P*_*r*_ as the system lingers for longer in each metastable state. At *τ* * however, the dynamics becomes effectively overdamped, and the time scales of *P* and *P*_*r*_ coincide as the system is macroscopically reversible. Dashed horizontal line represents half the mean first passage time estimated from the data as in Fig. 2(C). Error bars represents 95% confidence intervals over 200 non-overlapping trajectory segments of length 50000 s. (B) Lorenz system. The time scales from *P*_*c*_ are shorter than the reversibilized time scales of Fig. S3(A) across *τ*, reflecting the dissipative irreversible nature of the dynamics. Nonetheless, choosing *τ* * ≳ 0.1 s for robust almost invariant sets also yields nearly Markovian slow kinetics. In addition, the periodic structure of the Lorenz system is reflected in 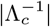, as it was for 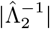, Fig. S3(A). Error bars represent 95% confidence intervals over 200 non-overlapping trajectories segments of duration 2000 s. (C) *C. elegans* foraging data. The inferred time scales of *P*_*c*_ reach a plateau after about *τ* = 0.75 s as in Fig. 4(C); dashed horizontal line highlights 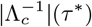. As in the Lorenz system, 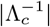 are shorter than 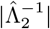 across *τ*, reflecting the large scale irreversibility of the dynamics. In fact, while 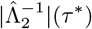 from *P*_*r*_ overestimates the expected 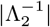 from “run” and “pirouette’ transition rates, the timescales obtained from 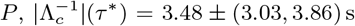, accurately predict the hopping dynamics. Error bars are 95% confidence intervals bootstrapped across worms.

**FIG. S10.**
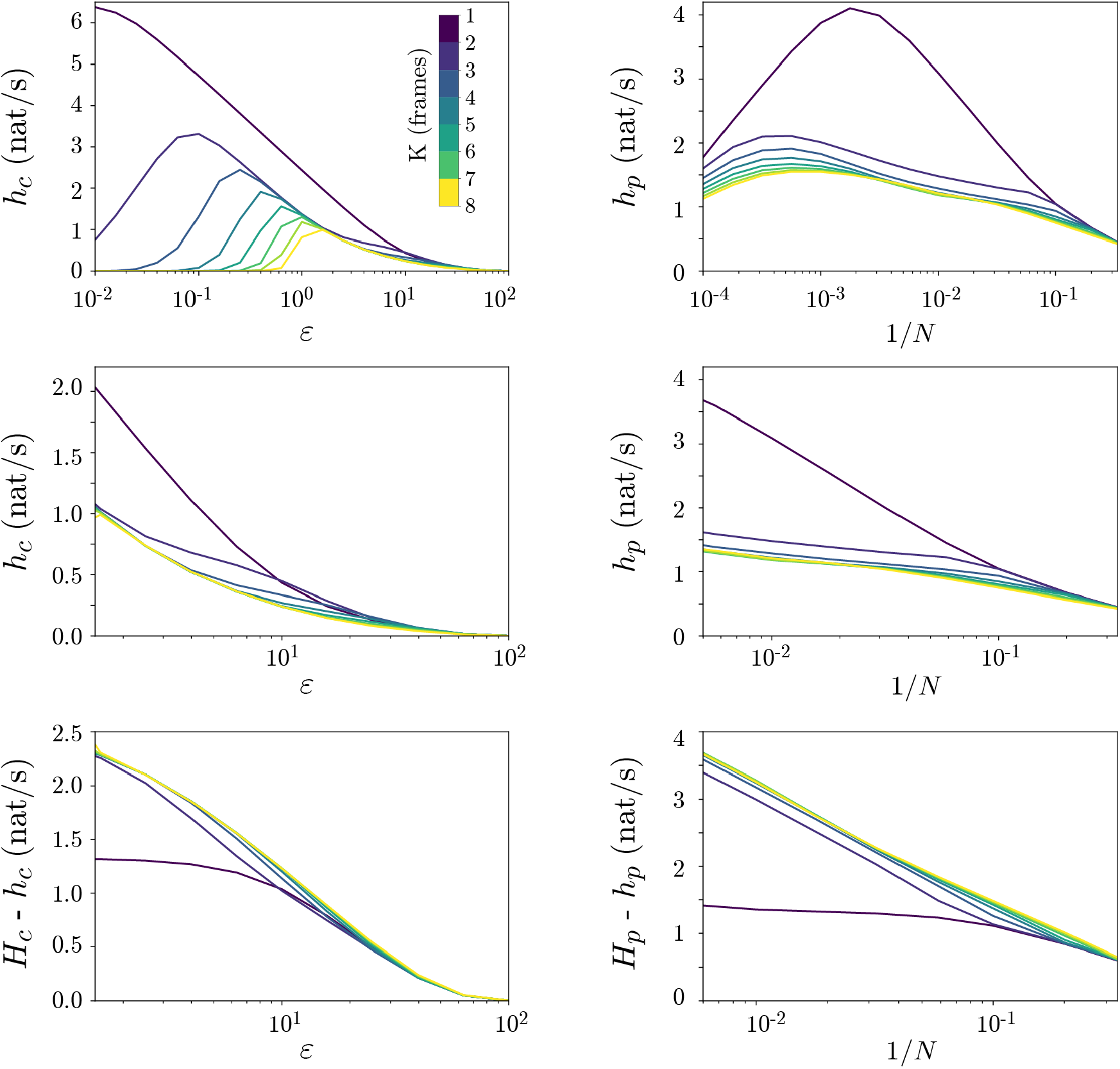
Comparison between the Cohen and Procaccia method [38] (left) and our partition based approach (right) to estimate entropy rates and predictive information as a function of the number of delays *K* for an AR(2) model. (Top) Entropy rate as a function of length scale *ϵ* computed using Eq. (18) (left) and as a function of the inverse of number of partitions 1*/N* computed using Eq. (4). Colors indicate the growing number of delays *K*. The geometric based approach suffers from finite size effects with the growing dimensionality of the space (from adding *K* into *y*^*K*^), but the entropy rate is generally higher for *K* = 1 than for *K* > 1. Similarly, *h*_*p*_ exhibits finite size effects for large numbers of partitions, and the estimates of the entropy rate drop. (Middle) Same as the top plot but now focused on a length scale regime at which the finite size effects of the estimates of entropy are not present. Note that the growth of the entropy with decreasing length scale (increasing number of partitions) is much fast for *K* = 1. (Bottom) Predictive information as a function of length scale *ϵ* (left) and the inverse of the number of partitions 1*/N* (right). For *K* > 1 the reconstructed state space maximizes the predictability.

## Footnotes

1 For simplicity, we use a slight abuse of notation: unlike the previous *π* from 𝒫_*τ*_, here *π* ∈ ℝ ^*N*^ is the finite approximation of the continuous invariant density.

2 We note that this is not a sufficient condition for Markovianity, as also the eigenvectors need to be constant with *τ*.

3 We note that on longer time scales the MSD exhibits the behavior of a confined random walk due to the rigid boundaries of the agar plate, which makes it non-trivial to accurately estimate the diffusion coefficient [81]. Nonetheless, the estimated effective diffusion coefficients provide a measure of the extent to which worms explore the environment.

